# Spatial transcriptomics identifies novel *Pseudomonas aeruginosa* virulence factors

**DOI:** 10.1101/2024.06.20.599896

**Authors:** Hao Zhou, Oscar Negrón, Serena Abbondante, Michaela Marshall, Brandon Jones, Edison Ong, Nicole Chumbler, Christopher Tunkey, Groves Dixon, Haining Lin, Obadiah Plante, Eric Pearlman, Mihaela Gadjeva

## Abstract

To holistically unravel the complexity of pathogen-host interactions within infected tissues we leverage a dual spatial transcriptomic approach that, for the first time, simultaneously captures the expression of *Pseudomonas aeruginosa* genes alongside the entire host transcriptome in a model of ocular infection. This innovative method reveals differential pathogen and host-specific gene expression patterns across specific anatomical regions generating a unified transcriptional map of infection. By integrating these data, we developed a predictive ridge regression model trained on images from infected tissues. The model achieved an R² score of 0.923 in predicting bacterial burden distributions by using host features thereby predicting novel biomarkers associated with disease severity. Our analysis revealed a complex interplay between *P. aeruginosa* nutritional requirements and protective host responses and identified novel interactions between bacterial metabolite transport proteins and host autophagy. Among an array of iron acquisition gene transcripts that showed significant enrichment at the host-pathogen interface, we discovered a novel virulence mediator PA2590. This study highlights the power of spatial transcriptomics, particularly in combining bacterial and host transcriptomes, to uncover novel host-pathogen interactions, advance our understanding of bacterial virulence mechanisms, and point to druggable molecules.

## Introduction

Genomic and transcriptomic methodologies including next-generation sequencing and high-throughput RNA analysis open possibilities for probing the intricate interactions between pathogens and their hosts. Traditionally, these approaches produce aggregate data from entire infected tissues (1–3). Missing from these analyses is spatial resolution, which can enhance our understanding of the bacterial and host response in infected tissues. We hypothesize that transcriptional analysis combined with spatial resolution will allow us to holistically explore how pathogens adapt and survive within specific tissue microenvironments. This approach also identified novel virulence factors that are potential targets for therapeutic intervention.

Data integration to achieve a spatially resolved understanding of infection remains a formidable challenge. To address this issue, our study employs dual spatial transcriptomics profiling, a novel approach that captures the expression profiles of bacterial gene transcripts alongside the complete host transcriptome. This methodology was applied to examine the transcriptional landscape of *P. aeruginosa*-induced infection in a model of acute ocular disease. By mapping the pathogen-specific transcriptional profiles at a localized scale, we reveal temporal site-specific transcriptional adaptations and novel virulence factors. To the best of our knowledge, this is the first example of the utility of spatial data to discover novel bacterial virulence mechanisms in a non-biased manner.

Previously, machine learning methods were used to diagnose viral and bacterial infectious diseases (4),(5) and predict the risk of secondary bacterial infections (6). Most of those analyses were performed on data acquired through microarray or RNAseq approaches that lack spatial resolution. In contrast to these studies, our spatial transcriptome data analysis leverages host transcriptome features to predict infection burden and disease severity. This analysis also correlates host transcript abundances with bacterial transcripts, thereby generating a unique tool to predict bacterial - host interactions.

Cumulatively, our findings not only highlight the significance of integrating spatially resolved transcriptomic data to explore the complexities of host-pathogen interactions, but also suggest potential avenues for the development of targeted therapeutic interventions.

## Results

### Spatial transcriptomics reveals tissue-segment specific host transcript enrichment

To generate a spatial transcriptional profile of infection, we used a well-characterized murine model of *P. aeruginosa* corneal infection. We utilized the Nanostring GeoMx platform for non-biased characterization of the host transcriptome (Fig. 1A). Sections from paraffin-embedded eyes were processed and we examined the responses in the corneal epithelium, the corneal stroma, and the anterior chamber as regions of interest (ROIs). Workflow is shown in Fig. 1A.

**Figure 1.**
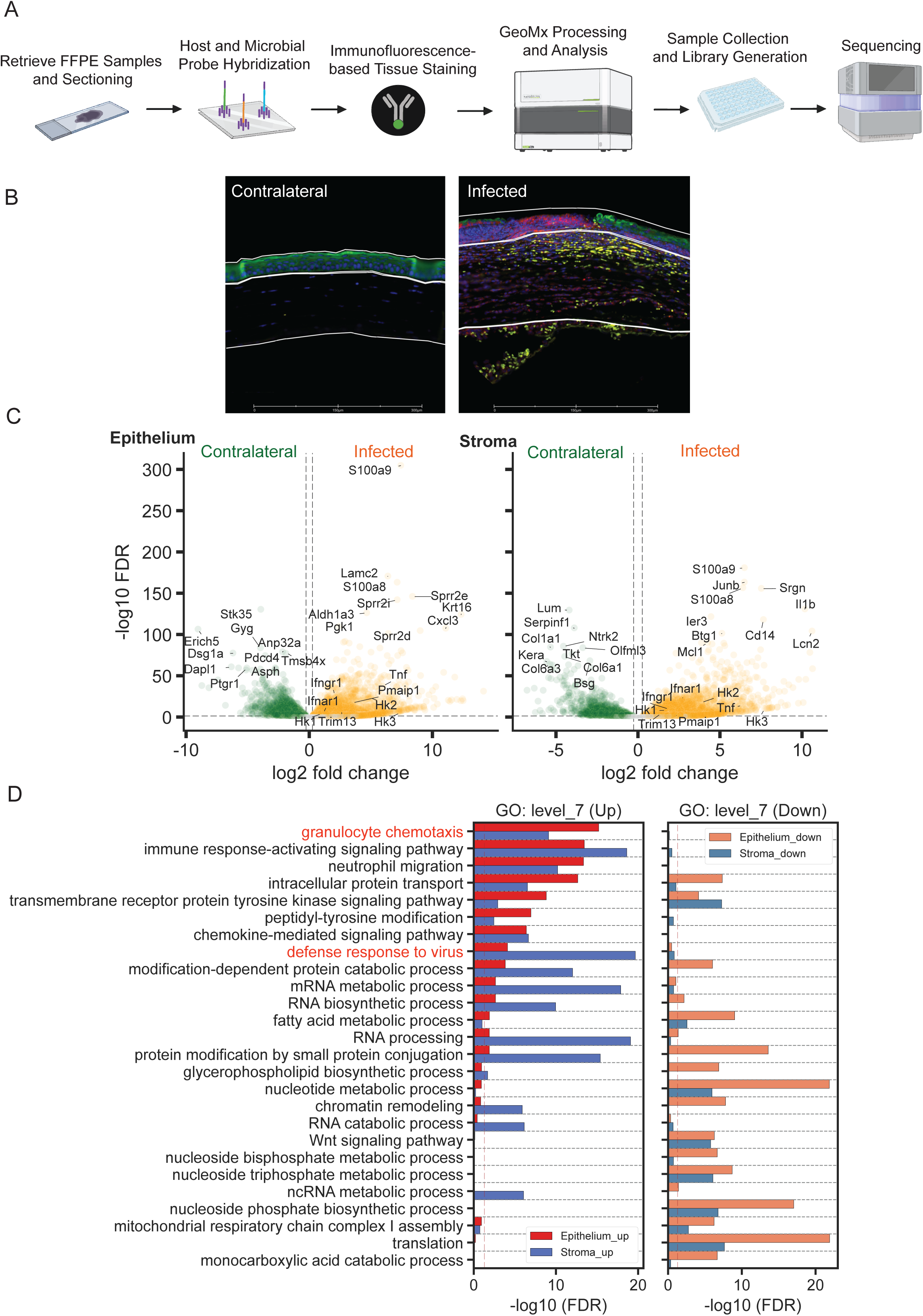
Non-biased whole mouse transcriptome analysis of P. aeruginosa-infected tissues reveal differential enrichment for host transcripts depending on the anatomical site. A. A schematic diagram of the workflow. Ocular tissues were harvested at 24h post-challenge with 1x10^6^ CFU PAO1. Tissues were paraffin-embedded, sectioned, stained for morphological markers and hybridized with whole host transcriptome probe library. Regions of interest were selected to encompass epithelial layer, stromal layer, and anterior chamber of the eye. Probes were cleaved post-hybridization, collected with GeoMx, amplified, and generated libraries sequenced. Image generated with BioRender. B. Representative immunohistochemical images of noninfected (contralateral, control eyes) and P. aeruginosa-infected ocular tissues stained for cytokeratin (green), Ly6G (red), DNA (blue), and c-KIT (yellow) were visualized with GeoMx (300 μm size bar, N=2 biological replicas). Robust neu- trophil infiltration (red) in the corneal stroma and epithelium is readily observed in the infected site. C. Volcano plots illustrating differentially expressed transcripts in epithelium and stroma tissue layers of the eye following infection, compared to tissues of the non-infected contralateral eye at 24 hours post-challenge. (N=2, biological replicas). The x-axis indicates fold change (log_2_ scale), and the y-axis shows the statistical significance (-log_10_ FDR-adjusted p-values). Sample metadata included. D. Bar plot of GO (Gene Ontology) functional enrichment analysis, highlighting tissue site-specific gene transcript clustering at level 7. The x-axis displays the FDR-adjusted p-values (-log_10_ scale), indicat- ing the statistical significance of the enrichment. The y-axis lists the level 7 GO categories with enriched transcripts. This analysis incorporates data from ROI profiles comprising 24 epithelial and 24 stromal samples from ocular tissues harvested at 24 hours post-challenge and 8 epithelial and 8 stromal samples from control samples. (N=2, biological replicas). GO enrichment analysis listed in Suppl. Table 2. **Cumulatively these data offer a comprehensive tissue map of infection indicating region-specific alterations.**

Corneal epithelial cells were identified as keratin 6^+^ve; neutrophils were identified as Ly6G^+^ve. Uninfected (contralateral eyes) and infected ocular tissues were examined (Fig. 1B). Transcriptome analysis revealed common and segment-specific enrichment of host and bacterial transcripts. Consistent with the rich neutrophil infiltration into the infected tissues, neutrophil-specific S100a8 transcripts were highly upregulated in the epithelium and stroma (Fig. 1C). Also upregulated were neutrophil and monocyte chemokines Cxcl2, and Ccl3, pro-inflammatory cytokines Il-1a and Il-1b, and the neutrophil iron chelating gene Lcn2. Cxcl2, Ccl3, Csf3, Cxcr4, and Il-1b, Il-1a, Il-12a cytokine transcripts were higher in the epithelium relative to the stroma (Fig. 1C). These data were reflected in the Gene Ontology (GO) enrichment analysis showing stronger granulocyte chemotaxis enrichment in epithelium versus stroma.

In contrast, transcripts associated with RNA metabolism, processing, ncRNA processing, and chromatin remodeling were predominantly upregulated in the stromal layer when compared to epithelium. Surprisingly, the infected stroma showed elevated anti-viral responses (Fig. 1D), including Type I IFN, IFNAR, and others (Suppl. Table 1).

After 48 hours of infection rapid disease progression was observed, as illustrated by elevated corneal opacity of the infected corneas (Suppl. Fig. 1A and B). The epithelial layer displayed enrichment of transcripts associated with metabolic alterations including glucose catabolism and cellular responses to glucose starvation, translation, and RNA processing (Suppl. Fig. 1D and Suppl. Table 2). In the stroma, we found sustained enrichment of transcripts associated with neutrophil recruitment (chemotaxis and migration), glucose catabolism, pyruvate metabolism and apoptosis (Suppl. Fig. 1C and Suppl. Table 2). Cumulatively, our data provide a temporal transcriptional and spatial record of tissue alterations during infection that are associated with neutrophil cytokine production and metabolic changes.

### Spatial analysis of bacterial transcript abundance in infected corneas reveals distinct enrichment profiles between the epithelium and stroma

Probes were generated for 100 *P. aeruginosa* genes that were encoded outer membrane proteins, extracellular proteins, reported virulence factors, and unknown, predicted genes. Probe development was based on the criteria of ORF frequency in >11,000 clinical isolates, evolutionary conservation, and their relation to known virulence. Three different gene-specific probes were designed for each of the bacterial genes of interest and used simultaneously to identify bacterial transcripts in the infected tissues (Suppl. Table 3).

Transcript expression was examined in PAO1 infected eyes in the positive corneal epithelium compared with the stroma and anterior chamber (AC). Representative images showed that infection caused loss of the cytokeratin staining in some of the surface layers of the cornea, consistent with the corneal perforation (ulceration) that occurs following infection. The underlying stromal layers showed rich Ly6G^+^ staining, indicative of neutrophil infiltration to infected cornea (Fig. 2B). Overall, we could distinguish ROIs with high and low bacterial transcript presence. This was reflective of the differences in the number of bacteria at distinct sites in the infected eye. While 97 transcripts were detected in almost half of all examined surface segments, only 37 transcripts were detected in most of the stromal segments.

**Figure 2.**
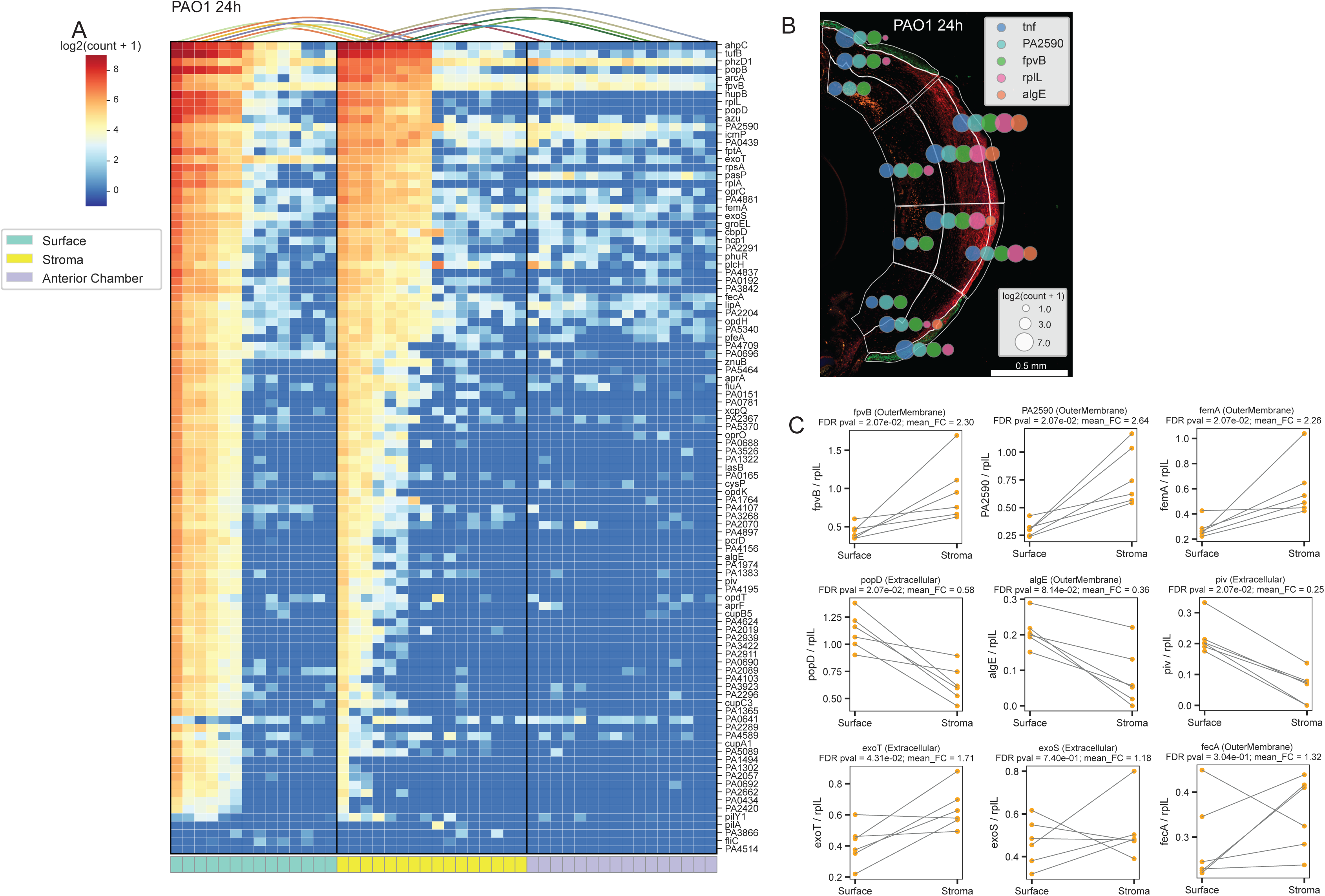
Bacterial transcript profiling of infected with P. aeruginosa PAO1 corneas reveal differential tissue enrichment. A. Heatmap showing the distribution of 100 bacterial transcripts in infected eyes. Each column represents data from individual ROIs. Surface epithelial ROIs (N=14) (green), stromal ROIs (16) (yellow), and Anterior Chamber (AC) ROIs (N=16) (purple) are grouped together. The curved lines on the top of the heatmap connect juxtaposing ROIs. Data are presented cumulatively from ROI selections derived from two biological replicas. B. Immunohistochemistry analysis of infected corneal section stained for cytokeratin (green) and Ly6G+ neutrophils (red). Size bar: 0.5 mm. Over- lay plot of host and bacterial transcript levels in PAO1-infected tissues. Logarithmic transcript abundances (log_2_(count + 1)) of host TNFα, and bacterial PA2590, FpvB, RplL, and AlgE are represented as circles, whose diameter reflects transcript burden in the individual ROIs. C. Bacterial enrichment was calculated as relative transcript abundance in the surface segments compared to the stromal segments. Bacterial transcript levels were normalized to the housekeeping bacterial RplL transcripts. FDR values and fold change (FC) are annotated per each gene. Representative bacterial enrichment data depicting: 1) transcripts that were enriched in the stroma FpvB, PA2590, FemA; 2) transcripts showing no enrichment, and 3) transcripts showing decreased abundances such as AlgE and Piv. Complete enrichment analysis is provided in Suppl Table 4. Data are representative of two experiments (Suppl. Fig. 2)

Interestingly, nine transcripts were consistently detected in the cytokeratin^+^ and cytokeratin^-^ surface layers, Ly6G^+^ and Ly6G^-^ stromal regions, and AC segments, including AhpC, TufB, PhzD1, ArcA, FpvB, PA2590, IcmP, LipA, and FemA (Fig. 2A and B).

Bacterial transcript enrichment analysis, normalized to the housekeeping RplL transcript abundance, reflected transcript increases in the infected stromal segments when compared to the infected surface layers (Fig. 2C). Nineteen transcripts showed significant upregulation in the infected tissues (Suppl. Table 4). Those included transcripts with known functions such as ArcA, PhzD1, IcmP, OprC, AhpC, LipA, ExoT, PhuR, OpdH, FpvB, and FemA, and transcripts encoding proteins that are predicted but have not been characterized: PA2590, PA0439, PA2204, PA2291, and PA4881. In contrast, transcripts for AlgE, PopD, and Piv were decreased in the stroma, while ExoS and FecA showed no enrichment (Fig. 2C and Suppl. Table 4).

To determine if bacterial transcript tissue penetrance is strain-specific, corneas were infected with the cytotoxic (ExoU expressing) PA14 strain. No significant enrichment of bacterial transcripts was observed at 24h post-challenge (data not shown); however, at 48h post-challenge, deeper bacterial transcript penetrance and significant changes were detected. Fourteen transcripts were similarly enriched between the PA14 and PAO1 data sets. These included PhzD1, FpvB, PA2590, IcmP, ArcA, FemA, LipA (Fig. 3B, C, and D). The transcriptional heat map comparisons between the sham and infected tissues confirmed the specificity of bacterial transcript detection (Fig. 3A and Suppl. Table 5). Cumulatively, these results highlight the appearance of bacterial transcriptional patterns in the infected tissues irrespective of the strain types and thereby illustrate spatial organization during infection.

**Figure 3.**
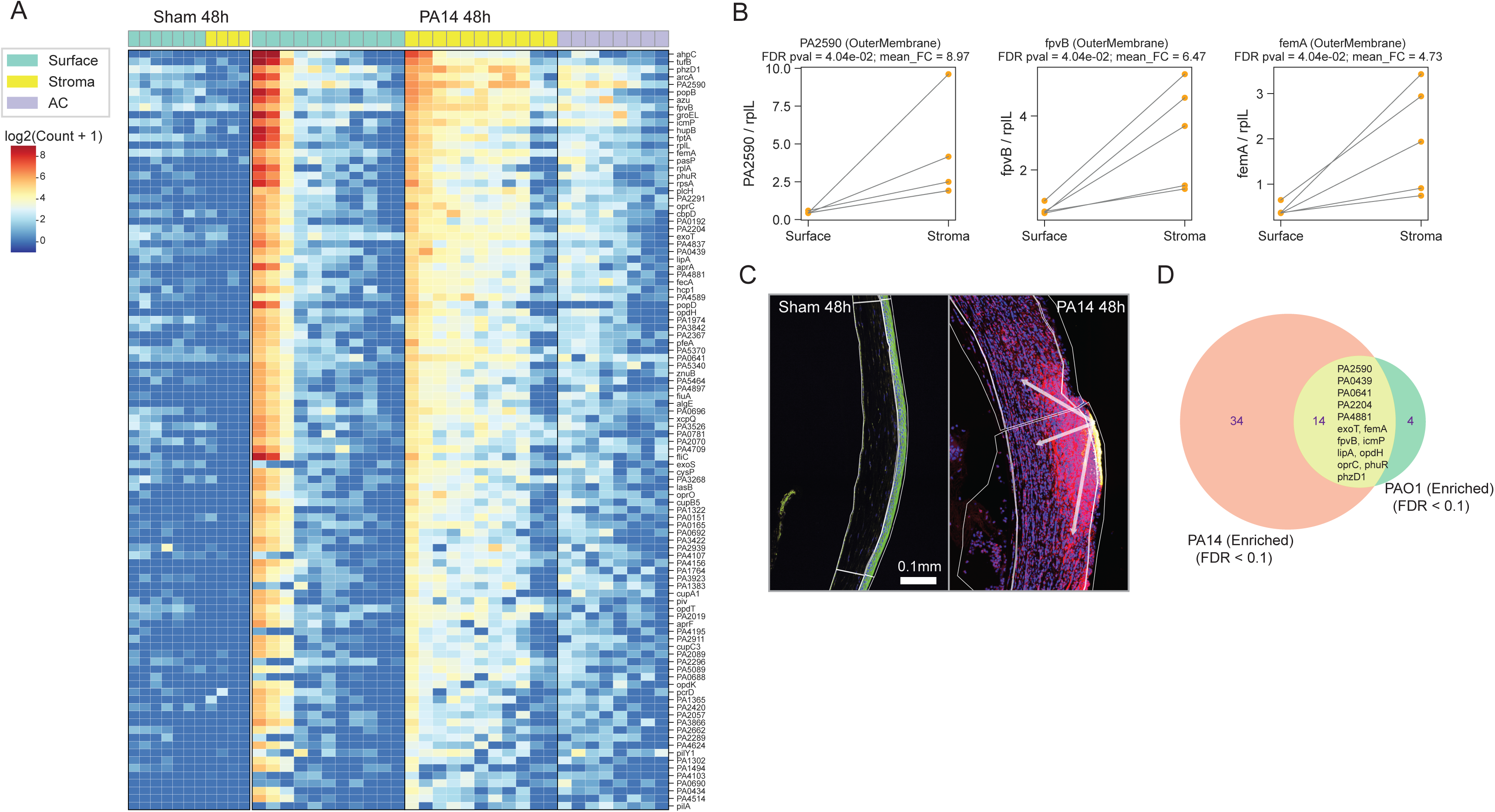
Bacterial transcript profiling of PA14-infected tissues exhibits similar tendencies as the PAO1-induced infections. A. Heat map showing the distribution of 100 bacterial transcripts in shams and PA14-infected eyes. The shams and the infected tissues were harvested and processed 48h post-challenge. Each column represents data from individual ROIs. Transcript abundances were compared between surface epithelial sham ROIs (N=7) (green), stromal sham ROIs (N=4) (yellow), and infected surface ROIs (N=11) (green), infected stromal ROIs (N=11) (yellow), infected AC ROIs (N=8) (purple) are grouped together. Data are presented cumulatively from ROI selections derived from three biological replicates. B. Bacterial enrichment was calculated as relative transcript abundances in the surface segments compared to the stromal segments. Bacterial transcript levels were normalized to the housekeeping bacterial RplL transcripts. FDR values and fold change (FC) are annotated for FpvB, PA2590, and FemA. Complete enrichment analysis is provided in Suppl Table 5. C. Immunohistochemistry analysis of shams and PA14-infected corneal section stained for OprI (yellow), cytokeratin (green), Ly6G+ neutrophils (red), and DNA (blue). Size bar: 0.1 mm. Arrows indicate bacterial spread from the infected surface lesion into the stroma. Shams, N=3, infected tissues N=11 biological replicates. D. Venn diagram depicting differentially present and shared bacterial transcripts that are enriched within the PAO1 and PA14-infected tissues. **Cumulatively, data point to commonalities in infection architecture that are not strain-specific.**

### Harnessing host-bacteria dynamics for accurate prediction of infection levels in tissue regions

Next, leveraging the host transcriptional data, we examined its functional relationship with the *P. aeruginosa* housekeeping gene RplL, which we used as a correlate for the number of bacteria in the infected tissues. Using Ridge Regression analysis (Fig. 4A) we demonstrated significant predictive accuracy, achieving an R² score of 0.923. The selection of 600 key host gene transcript features (Fig. 4B) refined the model, enhancing our understanding of the spatial distribution of bacterial burden across infection sites.

**Figure 4.**
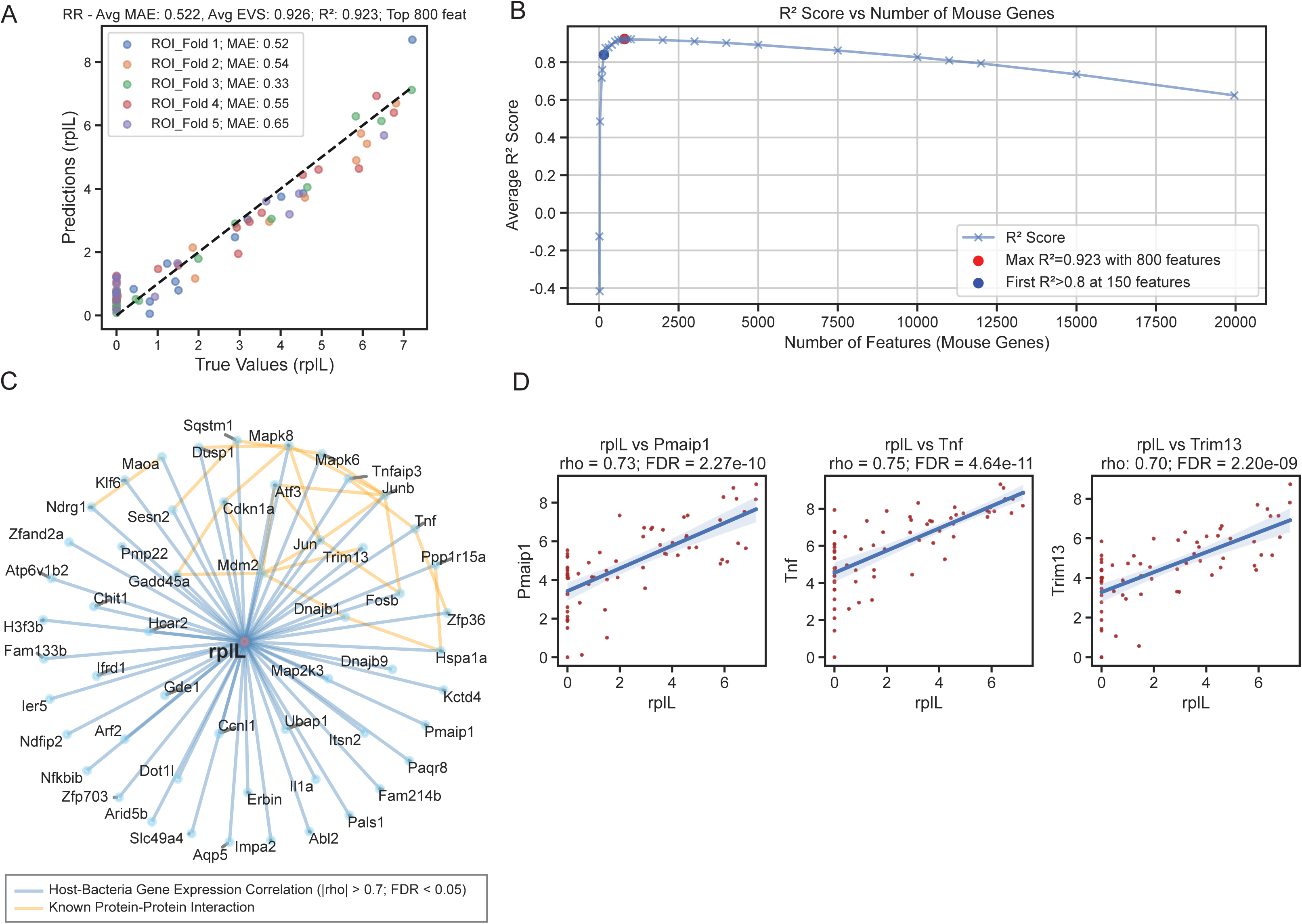
Harnessing host responses for highly predictive analysis of infection levels in tissue regions. A. Residual plot of a Ridge Regression model trained on host expression profiles to predict bacterial RplL gene expression levels. Data values are transformed using log_2_(values + 1). The model underwent five-fold cross-validation using ROIs from tissues derived from 24h post-challenge. Model performance was evaluated based on average MAE, EVS, and R2 scores. B. Evaluation of model performance when trained with varying numbers of mouse gene features. Top features were chosen for subset training based on the coefficient values from the model trained using the complete dataset. Average R^2^ scores provided the metric of assessment. Models, based on different feature sets, underwent optimization using grid search for hyperparameter tuning. C. Host-bacteria gene expression correlation network, including features with absolute rho values greater than 0.7 and an FDR less than 0.05, determined using Spearman’s correlation. **Cumulatively, these data offer a path towards predicting pathogen burden through the surrogate measurements of RplL abundance.**

We found that 150 host features were sufficient to accurately predict bacterial burden measured based on RplL transcript abundance, with an R² score of >0.8. We generated a host-bacterial RplL transcript correlation network and examined it for known host protein interactions (Fig. 4C). We found multiple signaling mediators associated with type I IFN, IL-1a, and TNF-driven inflammatory responses, which were highly interconnected for protein-protein interactions, further indicating the observation that the identified host response network reflected highly coordinated immune mechanisms (Fig. 4C). Examples for individual strong correlations between *P. aeruginosa* RplL and TNF-α, Trim13, and phorbol-12-myristate-13-acetate-induced protein1 (Pmaip1) are shown (Fig. 4D) and demonstrate that this approach can identify known (e.g., Il-1α, TNF-α) and unexpected immune determinants of infection including Trim13 and Pmaip1. Employing ‘leave-one-out’ validation on our model with a set of mouse hosts infected by PAO1 or PA14 strain, as illustrated in Suppl. Figure 2, showcases its adaptability and potential for broad applicability. This approach involves training the model on data from all but one host and assessing its predictive accuracy on the excluded host. The successful application of this technique across biological replicates demonstrates the model’s robust predictive capacity.

Cumulatively, our approach offers a detailed perspective on the variability of bacterial load across tissue regions and demonstrated the ability to predict bacterial tissue burden by select host transcript feature abundance, contributing to the field of precision medicine in diagnosing bacterial infections.

### Infection with the PA2590Tn deletion mutant strain is associated with reduced pathology and bacterial burden

Bacterial enrichment analysis identified known and unknown *P. aeruginosa* transcripts that were enriched in the infected tissues. To determine the functional importance of the predicted PA2590 gene that showed significant penetrance in the infected tissues from PAO1 and PA14-challenged mice, corneas were infected with *P. aeruginosa* PA14 WT parental strain or the PA2590 transposon mutant (PA2590Tn). Infected eyes were imaged, and corneal opacity, was imaged and quantified (Fig. 5). We also assessed viable bacteria by CFU. Since we established similar transcriptional enrichment for the PA2590 in the infected tissues of PAO1 and PA14- induced infections, infection studies were carried out with the PA14 strain.

**Figure 5.**
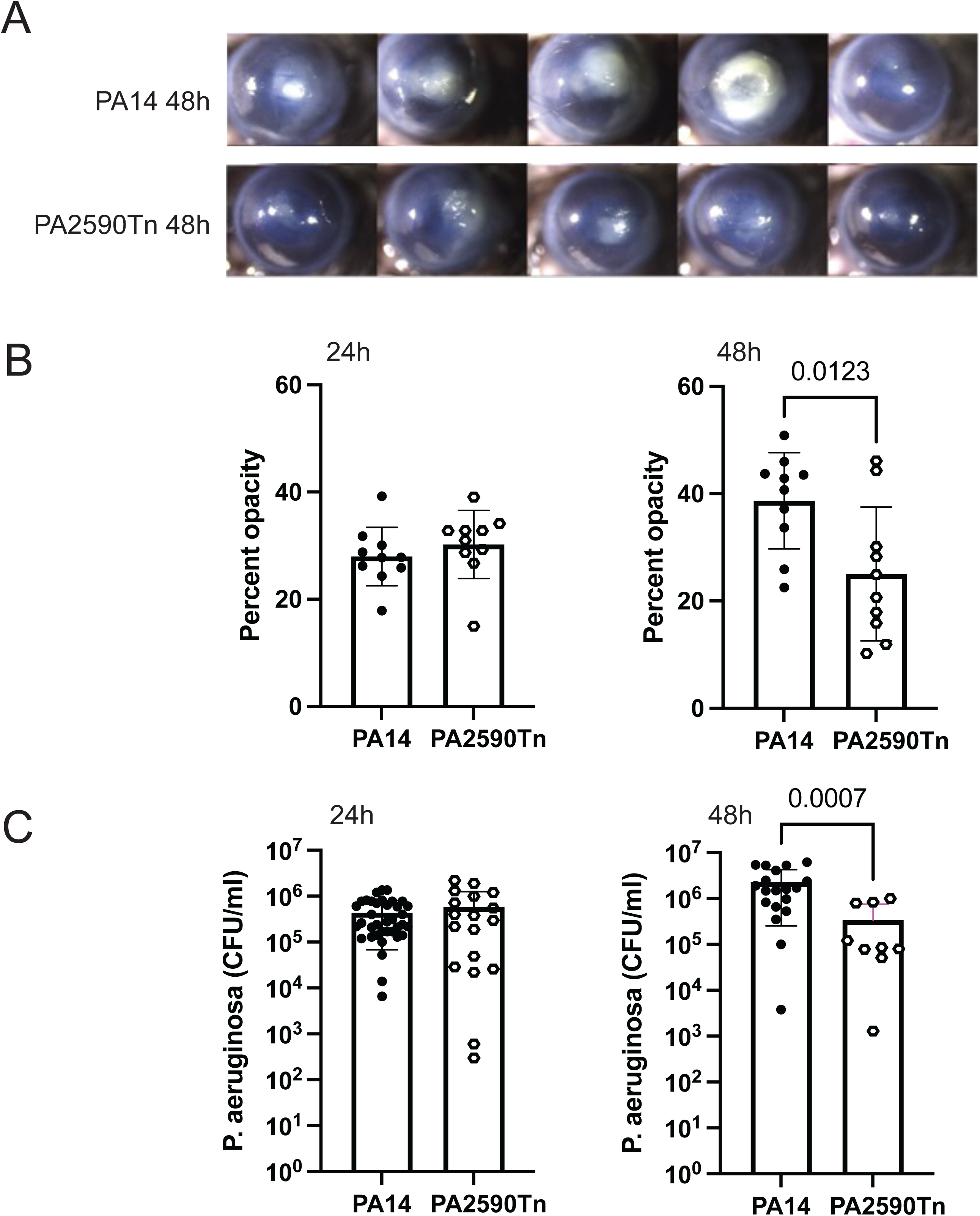
Infection with the PA2590Tn deletion mutant strain is associated with reduced pathology and bacterial burden. A. Representative images from infected with PA14 and PA14 PA2590Tn deletion mutant mice from two independent experiments. Images were collected at 48h post-challenge. B. Corneal opacity scores reflective of disease severity were quantified by image analysis at 24h and 48h post-challenge. Data are presented cumulatively from two independent experiments. The individual symbols present a single infected cornea. The filled symbols represent infected C57Bl6/J mice with the PA14 strain (N=10) and the open circles represent C57Bl6/J mice infected with the transposon deletion mutant for PA2590 (N=10). Unpaired Student’s t-test with Welch’s correction (P=0.01). C. C57Bl6/J mice were infected with 5x10^5^ CFU of PA14 (N=19) (filled circles) or the PA14 PA2590Tn deletion mutant (N=9) (open circles), and bacterial burden was quantified at 24h and 48h post-chal- lenge. Data are presented cumulatively from two independent experiments. The individual symbols present individual infected corneas. Unpaired Student’s t-test with Welch’s correction (P=0.0007). **Cumulatively, data point to decreased disease severity in the absence of the PA2590 gene product.**

There were no significant differences in corneal opacity or bacterial count at 24h post-challenge between the WT PA14 strain and PA2590Tn, consistent with the lack of enrichment for PA2590 at this data point (data not shown). However, the PA2590Tn mutant showed decreased virulence at 48h post-challenge as we recovered significantly less bacteria when compared to WT infected tissues (Fig. 5B, C, and D). As bacterial burden at 48h post-challenge is typically predictive of the disease outcomes in this model, we concluded that infection with the PA2590Tn mutant causes less severe disease.

### Structural comparison analysis suggests that PA2590 transports iron or vitamin B12

To understand how PA2590 might affect *P. aeruginosa* virulence, we examined its prevalence and conservation, and conducted structural studies. Analysis for PA2590 prevalence in 11,447 publicly available *P. aeruginosa* genomes derived from clinical isolates revealed that the PA2590 ORF is widely distributed and present in at least 97% of all clinical isolates (Fig. 6A). There were no significant differences in the PA2590 distribution based on geographical location of the isolates. Amino acid conservation analysis was also performed using a library of genomes derived from clinical isolates from different clinical presentations such as wound, blood, respiratory, eye, and genitourinary tract infections. A high degree of conservation was identified in all isolates at the amino acid stretches encoding the extracellular loops of the predicted structure of PA2590 with AlphaFold2 (Fig. 6B) (7). Cumulatively, these data suggested an important biological function for PA2590.

**Figure 6.**
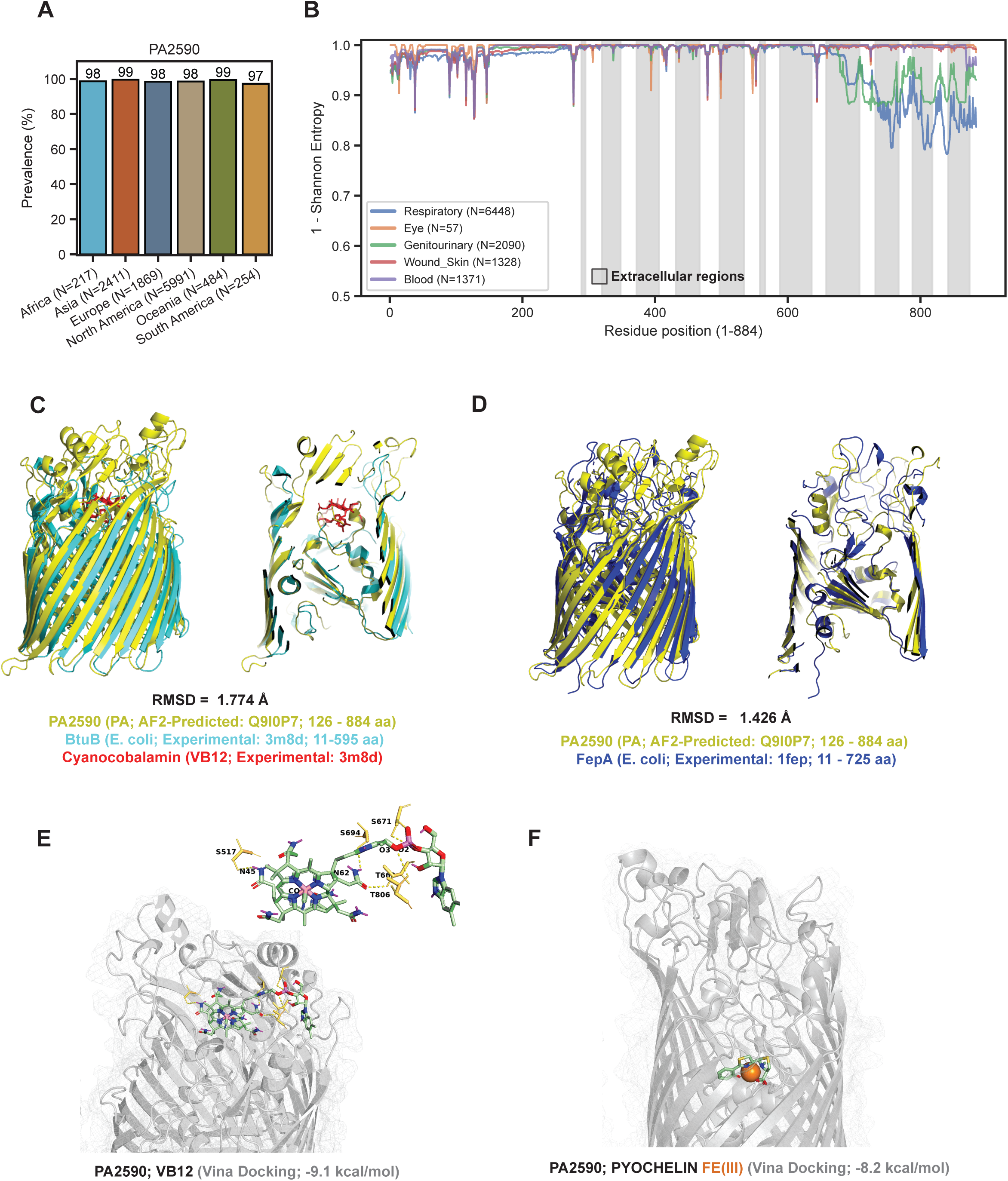
Structural homology-analysis predicts PA2590 gene to function as a vitamin B12 and iron transporter. A. Prevalence of the PA2590 gene in the genomes of P. aeruginosa clinical isolates sourced from various regions: Africa (N=216), Asia (N=2278), Europe (N=1680), North America (N=5763), Oceania (N=366), and South America (N=253). Here, ‘N’ denotes the total number of clinical isolates analyzed from each region. B. Amino acid conservation analysis for the PA2590 gene, illustrating data from clinical isolates of P. aeruginosa obtained from different infection types: genitourinary (blue line, N=1991), respiratory (orange line, N=6132), blood (green line, N=1202), and wound/skin infections (red line, N=1302). The x-axis displays the residue position, while the y-axis shows the 1-Shannon entropy values. Regions shaded indicate the extracellular loops of the protein. C. Structural homology modeling highlights parallels between the PA2590 structure predicted by AlphaFold2 and the experimentally determined structure of the E. coli BtuB protein. The figure contrasts the predicted model (light green) with the Btub structure (dark green), with VB12 depicted in red. Structural superimposition was conducted using the cmd.align function in PyMOL, involving 1000 alignment cycles with the transform parameter set to 1. This analysis resulted in a Root Mean Square Deviation (RMSD) of 1.774 Å for residues 126-884 of PA2590 and 11-595 of BtuB, indicating a significant level of structural similarity. D. Structural homology modeling highlights parallels between the PA2590 structure predicted by AlphaFold2 and the experimentally determined structure of the E. coli FepA protein. The figure depicts the predicted model (light green) with the FepA structure (dark blue). The RMSD is 1.426 Å over amino acid residues 126-884 of PA2590 and 11-725 of FepA, indicating a significant degree of structural similarity. E. In silico docking experiments were conducted between PA2590 and VB12 using Autodock Vina. The pose with the lowest energy (kcal/mol) was chosen for representation. Pymol scripts were used to highlight residues potentially interacting with PA2590, focusing on polar contacts within a 3.5 Å range. F. In silico docking experiments were conducted between PA2590 and pyochelin using Autodock Vina. The pose with the lowest energy (kcal/mol) was chosen for representation, indicating a docking score of -8.2 kcal/mol. **Cumulatively, data indicate that the PA2590 might function as either an iron imported utilizing a host siderophore or VB12 importer.**

Comparative structural analysis established a high degree of conformational similarities between PA2590 and *the E. coli* BtuB protein (PDB: 3m8d; R-Value Free: 0.275) present in *E. coli* (Fig. 6C). BtuB is a vitamin B12 (VB12) transporter, suggesting that PA2590 might have a similar function. Docking experiments revealed that PA2590 is likely to bind VB12 (Fig. 6E, Vina docking: -9.1 kcal/mol). For comparison, the docking score for BtuB and VB12 was -8.2 kcal/mol. In addition to BtuB, our analysis identified structural homology between PA2590 and *E. coli* FepA (PDB: 1fep; R-Value Free: 0.282) (Fig. 6D). FepA is an iron and VB12 transporter (8). The acquisition of iron depends on ferric enterobactin binding to FepA where the transporter preferentially recognizes the catecholate part of the complex (9). Unlike *E. coli*, our docking experiments did not indicate a potential interaction between enterobactin and PA2590. However, mammalian siderophores can be synthesized by eukaryotic homologues to the *E. coli* enzyme generating enterobactin (10). Therefore, we examined the potential binding of PA2590 to an abundant host siderophore 2,5-dihydroxybenzoic acid (2,5-DHBA). The *in silico* docking experiments confirmed the predicted interaction (Suppl. Fig. 3C, Vina docking: -6.0 kcal/mol).

*P. aeruginosa* uses two main siderophores for iron uptake: pyoverdine and pyochelin (11). As both known receptors were upregulated in the infected tissues and to determine if there might be redundancy in iron uptake, the potential interaction of PA2590 with Fe(III)-pyochelin was examined. The experiments showed a significant docking score of -8.2 kcal/mol, indicative of a role for PA2590 in the internalization of iron complexed with *Pseudomonas*-derived siderophores (Fig. 6F). Cumulatively, our data indicate that PA2590 mediates both iron and VB12 transport.

## Discussion

Bacterial infections remain a major global health threat, including those caused by *Pseudomonas aeruginosa*, a highly adaptable pathogen notorious for its resistance to antibiotics and its ability to cause severe infections in multiple tissues (12,13), (14). Despite its clinical significance, our understanding of the molecular mechanisms that underlie *P. aeruginosa*’s ability to thrive remains incomplete (15,16). The complexity of host-pathogen interactions within infected tissues is a significant obstacle in developing effective therapies and vaccines (17,18).

Spatial transcriptomic profiling of infected tissues provides information on gene expression patterns at specific infection sites. This report is the first to use dual spatial RNA-seq, combining bacterial pathogen-specific and host transcriptome data with spatial resolution into a unified, multidimensional dataset. The experimental approach represents a significant advancement over previous studies that used either sequential analysis of eukaryotic and prokaryotic features or reported bacterial analysis based only on 16S RNA seq (19,20). Our data illuminate differentially enriched bacterial transcripts at the host-pathogen interface, thereby identifying novel pathoadaptative mechanisms. The spatially resolved data enable machine learning approaches to connect host transcriptional networks with bacterial profiles, thereby identifying key nodes for targeted therapies.

Our study uncovered a strong, predictive correlation between a network of interconnected host features and *P. aeruginosa* housekeeping transcript abundances (RplL) during infection.

Using at least 150 host features, we successfully predicted bacterial burden in each tissue segment with a significant degree of accuracy. The correlative analysis identified known and novel biomarkers of infection severity including IL-1α, TNF α, TRIM13, Pmaip1, and Chit1. *P. aeruginosa* is a potent inducer of IL-1α and TNFα cytokines that orchestrate neutrophil infiltration to the infected site (20,21,22). Loss of IL-1α, unlike the loss of IL-1β, promotes resistance to infection (23,24). Conversely, increases in IL-1α correlates with worse disease illustrating that the ML-enabled analysis correctly identified IL-1α as a biomarker for *P. aeruginosa* disease severity. Novel biomarkers included increases in Tripartite Motif Containing 13 (TRIM13) and Phorbol-12-Myristate-13-Acetate-Induced Protein 1 (Pmaip1) transcripts (26). These genes regulate autophagy and apoptosis (27), and while direct studies linking Trim13 and Pmaip1 with *P. aeruginosa*-induced infections are lacking, their involvement in autophagy and apoptosis suggest mechanistic connections worth exploring in the future.

We predicted that the bacterial transcriptional characteristics would show distinct features depending on anatomical location. Consistently, bacterial enrichment analysis exhibited a higher tissue penetrance of transcripts associated with iron acquisition. *P. aeruginosa* requires iron to detoxify reactive oxygen species (ROS), and nitric oxide (NO) to ensure pathogen survival. *P. aeruginosa* has two major siderophores - pyoverdine and pyochelin, and the latter is essential for acute infection (33). Transcripts for the pyoverdine receptor, FpvB, and the ferric mycobactin receptor, FemA, were significantly enriched during infection. In contrast, the pyochelin receptor, FptA, did not show significant tissue enrichment, suggesting that there are other transporters for pyochelin import. To this end, we observed a significant penetrance of the PA2590 transcript encoding a predicted protein. Structural analysis showed that PA2590 has a 22-stranded antiparallel β-barrel domain containing Ton B-dependent transporters and that 2590 has a high degree of structural similarity to *P. aeruginosa* FemA transporters, thereby indicating a possible role for PA2590 in mediating iron intake. Further docking analysis are indicative of pyochelin binding.

PA2590 is also structurally similar to dual iron and vitamin B12 BtuB transporter in *E. coli,* suggesting that PA2590 has a role in cobalbumin import (34). The dual import of iron and vitamin B12 reflects the need for *P. aeruginosa’*to survive in nutritionally poor, oxygen-limited environments (35). Interestingly, *P. aeruginosa* synthesizes vitamin B12 under aerobic conditions although this activity is suppressed under anaerobic conditions (36). The import system that facilitates anaerobic vitamin B12 intake is presently unknown (37)., and our analysis offers insights into mechanisms of vitamin B12 import. In *P. aeruginosa*, vitamin B12-dependent enzymes such as methylmalonyl-CoA mutase (38), methionine synthase (39), ethanolamine ammonia-lyase (40), glycerol dehydratase (41), and diol dehydratase (42) play crucial roles in metabolic versatility that demonstrates the ability of these bacteria to use alternative carbon sources. Vitamin B12 also supports DNA replication under anaerobic conditions by serving as a cofactor for essential ribonucleotide reductases (37). We also showed that PA2590 is functionally important in a clinically relevant model *P. aeruginosa* corneal infection as significantly less bacteria were recovered in the absence of PA2590. Cumulatively, these data demonstrate that this bioinformatic approach can identify novel pathoadaptation mechanisms, including bacterial virulence factors.

Collectively, our analyses highlight transcriptional changes associated with host autophagy pathway and metabolite acquisition by the pathogen, which indicates that there may be a causative link between bacterial vitamin B12, iron acquisition, and autophagy. Indeed, several reports show that iron sequestration by the host to limit bacterial infection promotes autophagy (43)(44)(45). Additionally, iron-siderophore complexes incubated with *C. elegans* induces mitophagy, a form of damaged mitochondrial autophagy (46). Conversely, inhibition of mitophagy renders *C. elegans* more susceptible to *P. aeruginosa*-induced infection (46). Vitamin B12 deficiency, similar to iron starvation, promotes autophagy due to reduction in the citric acid cycle and ATP production (47). Cumulatively, the spatially resolved data capture efficiently a complex host-pathogen interplay that identify key metabolites such as iron and vitamin B12 that regulate autophagy can also mediate resistance to infection.

In conclusion, the uses of spatially resolved maps of infection are widely applicable towards gaining foundational insights into host-bacterial pathogen interactions that will guide future mechanistic research. Our spatial transcriptome data analysis approach identified novel virulence mechanisms associated with key requirements for pathogen survival during infection.

## Methods

### Bacterial strains

*P. aeruginosa* strains PAO1, PA14, PA14_30590_Tn (PA2590 transposon mutant) were grown in Luria–Bertoni broth (LB) broth (BD-Difco) overnight at 37°C with agitation (200rpm). Cultures were then passaged in fresh LB broth (BD-Difco) until an OD600 of 0.20 (midlog phase) was reached. The PAO1 strain was obtained from Dr. A. Rietsch (Case Western Reserve University). The PA14 and PA14_2590_Tn were generated during the PA14 transposon insertion mutant library generation ((http://ausubellab.mgh.harvard.edu/cgi-bin/pa14/home.cgi)) and obtained from Prof. G. Pier (BWH, HMS) (48,49). The PA2590 transposon mutant was sequence validated (data not shown).

### Corneal infection model

Male C57BL/6 (“WT”) mice were purchased from The Jackson Laboratory (Bar Harbor, ME) and were housed in a B-level UCI vivarium room. Mice were maintained by the Pearlman Lab and UCI ULAR staff. Infected mice were transferred to D-level holding rooms until the experimental endpoint was reached. All protocols were approved by UCI Institutional Animal Care and Use Committee under IACUC protocol AUP-21-123.

Overnight cultures of *P. aeruginosa* were grown to an OD600 of 0.20 in LB broth, and cultures were then washed and resuspended in 1x sterile PBS (Corning). C57BL/6J mice were anesthetized with ketamine/xylazine (MWI Animal Health) and corneal epithelium was abraded with three, parallel 10mm scratches, performed with a 26 ½ -gauge needle (BD). *Pseudomonas* was then applied to the abraded cornea topically with 2µl containing 5x10^5^ CFU in PBS in the PA14 infection studies and with 5ml of 1x10^6^ CFU in the PAO1 infection studies. For use in the infection experiments, 100µl of overnight culture was added to 10ml of fresh LB and was subcultured for 2.5-3h to reach OD600=0.20. 1ml of culture was spun down at 3,000 x g (rotor, centrifuge) for 10 min and diluted to a desired inoculum. Experiments were carried out predominantly with male C57Bl6/J mice. After 24h or 48h, mice were euthanized, and corneas were imaged by brightfield microscopy to observe opacification. Bacterial CFU was quantified at 24h and 48h by serial dilutions of whole eye homogenates onto LB plates and were counted manually.

### Corneal opacity

Corneal images were taken using a Leica MZ10 F Modular Stereo Microscope with a Leica DFC450 C camera attachment. Image brightness was adjusted manually using ImageJ software (NIH) and was kept consistent for all images. Images were then converted to 8-bit for quantification. Using ImageJ, ROIs were drawn around the corneal area only (excluding the sclera) and mean intensity was recorded for each ROI. Images were corrected by subtracting intensity caused by glare of dissection lights (glare values were based on naïve corneas). Percent corneal opacity was calculated by dividing corrected mean intensity by the intensity of a completely white cornea.

### Bacterial quantification

Once images were taken, eyes were resected and placed into a 2 ml centrifuge tube containing 1ml of sterile 1x PBS and a sterile steel ball bearing (5 mm diameter). Tubes were placed in a Qiagen Tissue Lyser II and were homogenized at 30Hz for three minutes. After lysis, homogenates were serially diluted up to 1:10,000 and 10µl of each dilution was plated onto one LB agar (BD-Difco) plate (concentrations plated: undiluted, 1:10, 1:100, 1:1,000, 1:10,000).

Dilutions were streaked down the plate and plates were incubated at 37°C with 5% CO2 overnight and colony forming units (CFU) were counted manually the next day. CFU was then quantified as CFU/ml.

### Ocular tissue processing

After imaging the corneas, eyes had 15µl of 4% formalin (ThermoFisher) dropped onto them and were left to fix at room temperature for 15 minutes. Eyes were then carefully resected and placed in 1ml of iced 4% formalin (ThermoFisher) and were fixed at 4°C for 48-72hrs. After fixation, formalin was removed, and tissue was washed twice with sterile 1x PBS. Samples were then placed in 2ml of 70% ethanol and were kept at 4°C until shipment to UMass Histopathology was complete. Tissue embedding was done by UMass Histopathology. All tissues were processed using UMass Chan Medical School Morphology Core services following standard protocols.

### Target selection and probe designs

Probes targeting 100 selected genes in the PAO1 strain of *P. aeruginosa* were designed using Nanostring in-house scripts (Supplementary Table 1). These probes were designed to specifically bind to *Pseudomonas* genes, avoiding cross-reactivity with mouse genes and other microbes.

### Spatial transcriptomics workflow

5um thick sections from formalin-fixed paraffin embedded (FFPE) tissue blocks were mounted onto Superfrost Plus slides and prepped for GeoMx spatial transcriptomic profiling. Briefly, slides were baked at 37° C for 1 h prior to loading on a BOND RXm (Leica) for further baking, rehydration, heat-induced epitope retrieval (ER2 for 20 min at 100° C) and enzymatic digestion (0.1 ug/mL proteinase K for 15 min at 37° C). Tissue sections were then hybridized with the Nanostring mouse Whole Transcriptome Atlas (WTA) and custom *P. aeruginosa* probe sets at 37° overnight. Following hybridization, two, 25min stringency washes were performed to remove off-target probes and sections were subsequently blocked and stained with the following morphology marker antibodies: DNA (Syto13; 488 channel; Nanostring), PanCK (532 channel; Nanostring), OprI (Opr17_VHHFC; 594 channel; Thermo Fisher Scientific) and Ly6G (1A8; 647 channel; Novus Biologicals). Slides were then loaded into the GeoMx DSP instrument where the entire sections were scanned and imaged. Regions of interest (ROIs) were manually drawn on the section images and approved for collection. The GeoMx DSP then directed UV light to each ROI to cleave the barcodes (RNA ID and unique molecular identifier-containing oligonucleotide) from the bound probes. Illumina i5 and i7 dual-indexing primers (Illumina) were added to the collected barcodes during polymerase chain reaction (8 µl collected oligonucleotides/ROI) to uniquely index each ROI. AMPure XP beads (Beckman Coulter) were used for purification of PCR. Library concentration was measured using a Qubit fluorimeter (Thermo Fisher Scientific), and quality was assessed using a Tape Station (Agilent Technologies).

### Illumina Sequencing and Processing

The sequencing depth requirement was determined by aggregating the areas of illumination (measured in µm²) and multiplying the total area by a factor of 100, as recommended in the GeoMx DSP Library and Sequencing Guide (nanostring.com). Sequencing was executed using the NovaSeq 6000 System. The resultant FASTQ files were processed to Digital Count Conversion (DCC) files utilizing the GeoMxNGSPipeline v.2.3.3.10, a pipeline publicly available through NanoString Technologies. This pipeline facilitated individual probe alignment by identifying read tag sequences (RTS). Additionally, it eliminated duplicates in probe counts generated during the library preparation’s polymerase chain reaction, through a meticulous analysis of the unique molecular identifiers (UMIs) associated with each probe.

### Quantification and Statistical Analysis

The DCC files underwent analysis using NanoString Technologies’ GeoMx™ Digital Spatial Profiler. Key metrics such as the number of raw reads, the alignment rate of these reads, and the sequencing saturation for each area of illumination were scrutinized to evaluate the quality of library preparation. Criteria for inclusion in the analysis were set: more than 10^6^ raw reads per ROI, an alignment rate exceeding 80%, and a sequencing saturation greater than 80%. A "no template control" sample, devoid of target probes, was integrated into the library preparation and subsequent sequencing to gauge potential contamination. To estimate background noise and determine the limit of quantification, negative control probes employing External RNA Controls Consortium (ERCC) sequences were included in each GeoMx™ RNA assay. Target probe data were filtered based on frequency within each ROI, followed by normalization against the highest negative control counts using background subtraction. Differential gene expression analysis across various groups was performed using DESeq2, a statistical method designed to identify significant changes in normalized gene expression levels (50). The Python-based library GOATools were used for the functional enrichment analysis (51).

### Analysis of PA2590

To identify the PA2590 gene, we analyzed more than 11,000 genome sequences from clinical isolates collected between 2000 and 2023. These sequences, sourced from the NCBI database, were subjected to BlastP analysis with the criteria of >85% identity and >70% query and >70% subject coverage, to identify homologous genes of PA2590. Following this, Clustal Omega was utilized for sequence alignment (52). The conservation of amino acid residues within PA2590 was quantitatively evaluated using Shannon Entropy (53). This approach effectively highlights conserved regions by examining the variability of amino acid positions across the aligned sequences, offering insights into the evolutionary stability of the gene. MembraneFold was employed to predict membrane topology of PA2590 (54).

### Structural homology analysis and molecular docking

For structural insights, we employed Foldseek to align PA2590 (www.uniprot.org/uniprotkb/Q9I0P7/entry#structure) with existing structures in the Protein Data Bank (PDB) in 01/2024 (55). This alignment was instrumental in hypothesizing the functional structure of PA2590, based on similarities with known protein structures. Structural superimposition was conducted using the cmd.align function in PyMOL v2.5.4, involving 1000 alignment cycles with the transform parameter set to 1. *In silico* docking experiments were conducted using Autodock Vina (56). The pose with the lowest energy score (kcal/mol) was chosen for representation. Pymol scripts was used to highlight residues potentially interacting with PA2590, focusing on polar contacts within a 3.5Å range.

### Predictive Models and Correlative Analysis

We employed Random Forest (RF), Ridge Regression (RR), and Support Vector Machine (SVM) algorithms for predictive analysis, with these models trained on a complete set of host features (as detailed in Suppl. Fig. 2A). Among these, Ridge Regression emerged as the most accurate model and was selected for further in-depth analysis and optimization.

The Ridge Regression model was trained using host response profiles and subsequently used to predict the expression data of the PA RplL gene. For normalization of data values, a log transformation (log2(values + 1)) was applied. To assess the model’s predictive accuracy, we implemented a five-fold cross-validation strategy, utilizing Regions of Interest (ROIs) from mice that had been infected. Additionally, cross-validation was carried out across different hosts to validate the model’s capacity to predict outcomes in hosts at similar infection stages, as detailed in Supplemental Figure 2. The effectiveness of the model was thoroughly evaluated based on the average Mean Absolute Error (MAE), Explained Variance Score (EVS), and R² scores.

Further, we assessed the model’s performance when trained with varying numbers of mouse gene features. For this, we selected the top-performing features based on their coefficient values in the model trained with the complete dataset. The primary metric for assessing these subset-trained models was the average R² score. To optimize these models, based on different sets of features, we applied a grid search technique for hyperparameter tuning, ensuring that each model was fine-tuned for the best possible predictive performance.

We assessed correlations between bacterial RplL and host gene expressions using Spearman’s rank correlation, with p-value adjustments made through the Benjamini-Hochberg FDR method.

## Supplementary Tables

**Supplementary Table 1.** Host-specific transcriptional changes induced by infection at 24 hours post-challenge. This includes GO enrichment analysis results (Supplementary data to Figure 1D).

**Supplementary Table 2.** List of genes differentially expressed in infected epithelial and stromal ROIs when comparing 48-hour infection to 24-hour infection.

**Supplementary Table 3.** List of 100 *P. aeruginosa* transcripts profiled in the GeoMx experiments and corresponding Nanostring detection probes.

**Supplementary Table 4.** Enrichment analysis reflecting *P. aeruginosa* PAO1 bacterial transcript fold changes at 24h post-challenge.

**Supplementary Table 5.** Enrichment analysis reflecting *P. aeruginosa* PA14 bacterial transcript fold changes at 48h post-challenge.

## Data and Code Availability

Upon acceptance of the manuscript, the raw sequencing data from our genomics and spatial transcriptomics studies will be made available in the NCBI database. Additionally, the code used for training predictive models and conducting correlative analysis can be found at https://github.com/modernatx/spatial_RNAseq_analyses.

### Acknowledgments

We would like to acknowledge the outstanding contributions of the NanoString team: Maxine Macclain, Espy Anguiano, Tim Riordan, Ozge Cagsal-Getkin, Mary Hentschel, and Ankush Tyagi who worked tirelessly to help us conduct the feasibility studies and establish the technology. We would like to thank University of Massachusetts Histopathology core, Yu Liu and Jayme Haywosz, for embedding and processing services. We would like to thank Prof. G. Pier (MGB, HMS) for providing the PA14 and PA14 PA2590 strains.

## Author contributions

HZ conducted all computational analyses. SB and MM carried out all in vivo experiments and analyzed their results. ON was responsible for data acquisition within the GeoMx workflow. BJ primarily handled the sequencing analysis portion of the GeoMx workflow. EO and HZ selected the PA genes. CT advised on the sequencing aspects of the GeoMx workflow. GD and HL validated the PA2590 Tn mutant. HZ, NC, EP, and SB played key roles in study design and data interpretation. MG devised the study concept, and contributed to data collection, analysis, and interpretation. HZ, ON, NC, EP, and MG drafted the initial manuscript. All authors discussed the data, reviewed, and revised the manuscript.

## Ethics declarations

HZ, ON, NC, BJ, EO, CT, GD, HL, OP, and MG are employee of Moderna Inc. and may hold stock/stock options in Moderna, Inc. SA, MM, and EP have no conflicts to declare.

---

**Supplementary Figure 1.**
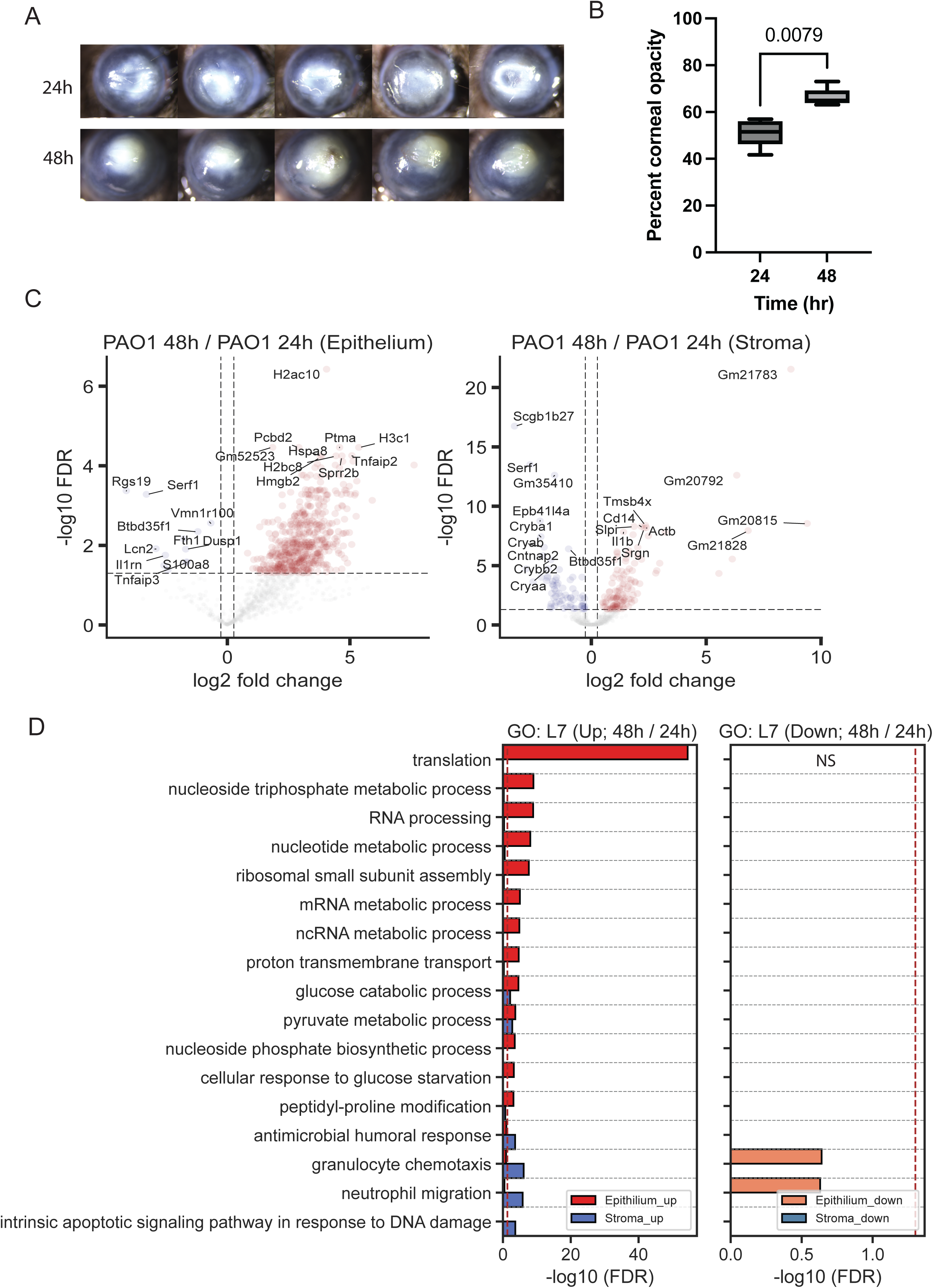
Temporal profiling of host response confirms sustained transcriptional remodeling and tissue-specific gene enrichment. A. Representative images from infected with 5x105 CFU P. aeruginosa PAO1 strain C57Bl6/J mice (N=5 biological replicas). Images are collected at 24h and 48h post-challenge. B. Corneal opacity scores reflective of disease severity were estimated at 24h and 48h post-challenge. (N=5 biological replicas). C. Volcano plots illustrating differentially expressed transcripts in epithelium and stroma tissue layers of the eye following infection, compared to tissues of the non-infected control eyes at 48h post-challenge. Mice were challenged with 5x105 CFU P. aeruginosa PAO1 and tissues were harvested at 24h (N=1) and 48h post-infection (N=2). Data are presented cumulatively. The x-axis indicates fold change (log2 scale), and the y-axis shows the statistical significance (-log10 FDR-ad- justed p-values). Detailed sample metadata is available in Suppl. Table 2. D. Bar plot of GO (Gene Ontology) functional enrichment analysis, highlighting tissue site-specific gene transcript clustering at level 7. The x-axis displays the FDR-adjusted p-values (-log10 scale), indicating the statistical significance of the enrichment. The y-axis lists the top level 7 GO categories with enriched transcripts. This analysis incorporates data from ROI profiles comprising 24 epithelial and 24 stromal samples from ocular tissues harvested at 24 hours post-challenge and eight epithelial and eight stromal samples from non-infected eyes, control samples.

**Supplementary Figure 2.**
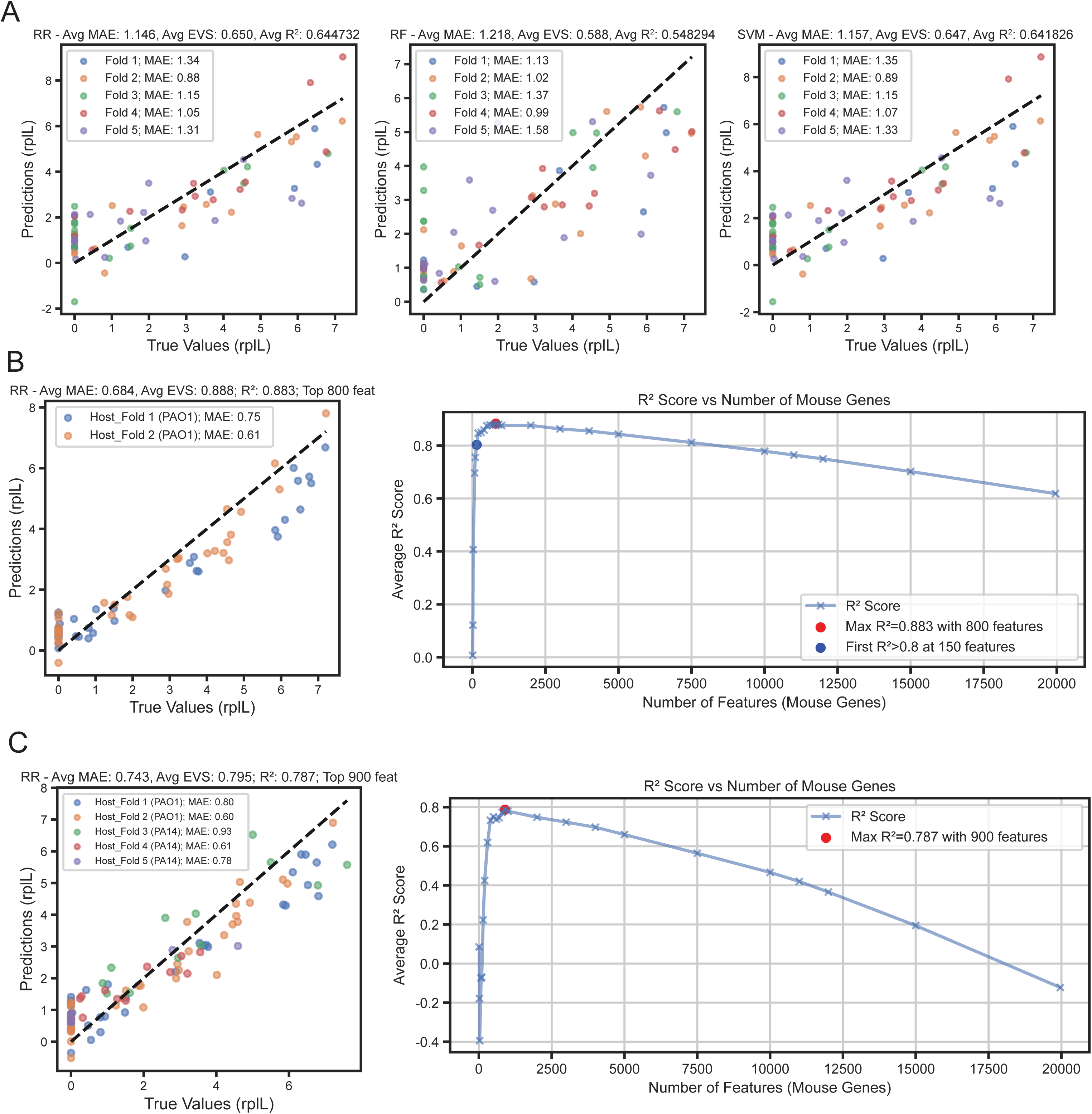
Evaluation of model performance in predicting pathogen-specific gene expression based on host response profiles from multiple biological replicas. A. Residual plots of models RF, RR, and SVM generated from host response profiles to predict patho- gen-specific RplL gene expression levels. Data values are log_2_-transformed (values + 1). Models were trained using ROIs selected from tissues collected 24h post-challenge and cross-validated against ROI datasets from different biological replicas. Model performance was assessed using average MAE, EVS, and R^2^ scores. B. Evaluation of model performance when trained on a single biological data set and validated against a second data set derived from a biological replica. Average R2 scores provided the metric of assessment. Models, based on different feature sets, underwent optimization using grid search for hyperparameter tuning. C. The assessment of model performance involved training on multiple biological replicas and validating on a single excluded replica, which included samples infected with PA14 and PAO1. Average R^2^ scores served as the evaluation metric. The models, based on various feature sets, underwent optimization through grid search for hyperparameter tuning.

**Supplementary Figure 3.**
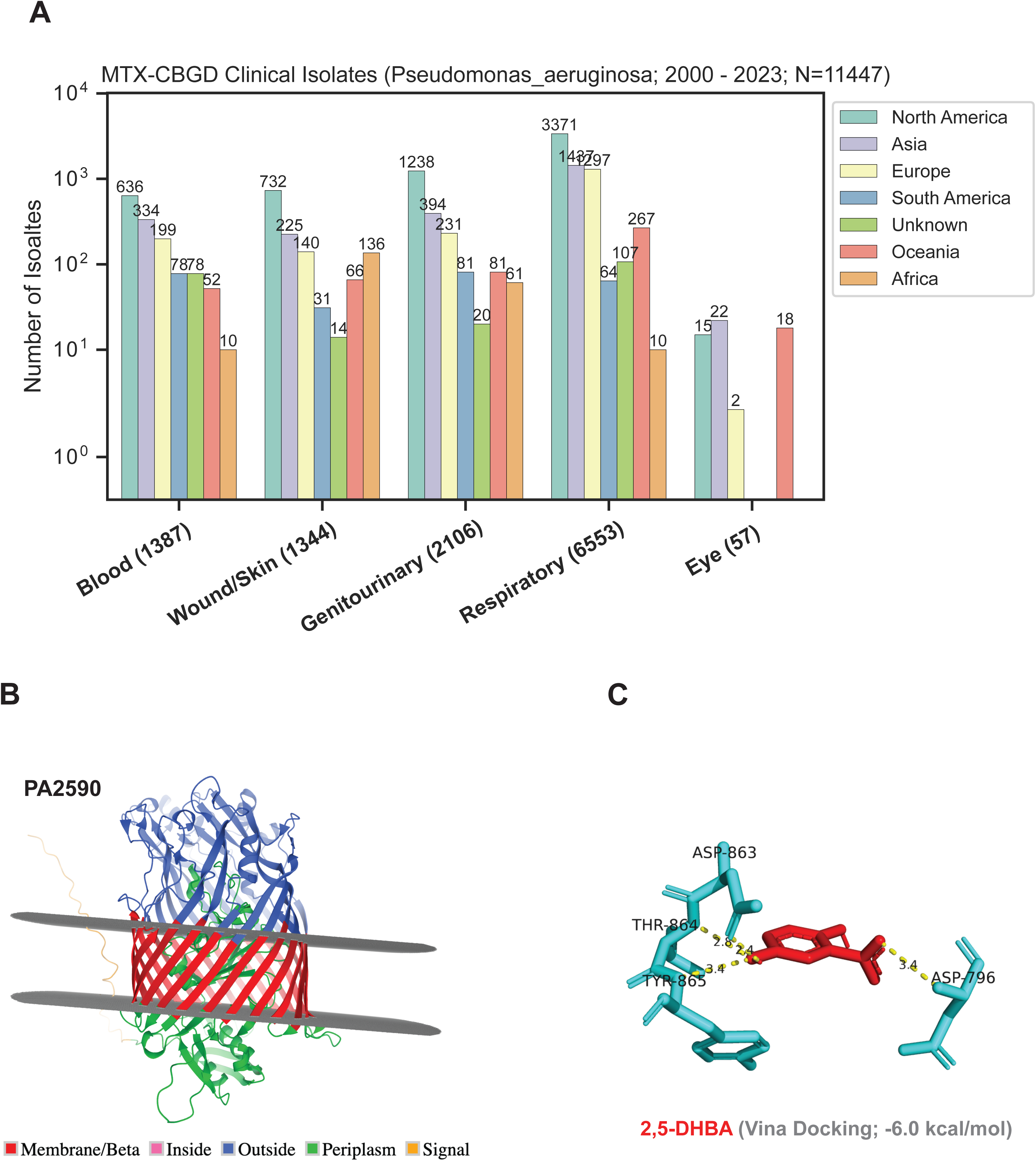
Structural and functional insights into PA2590. A. Genomic data from 11,447 *P. aeruginosa* clinical isolates was sourced from NCBI reflecting strain diversity for the past 23 years. The bar graph depicts the global distribution of the number of isolates plotted on the y-axis, and the types of associated infections plotted on the x-axis. B. Membrane topology of PA2590 structure predicted by MembraneFold. C. In silico docking experiments were conducted between PA2590 and 2,5-DHBA. The pose with the lowest energy (kcal/mol) was chosen for representation.

