## Supplementary material for "Spatial transcriptomics identifies novel *Pseudomonas aeruginosa* virulence factors": Supplementary Table 3.docx

| **Name** | **Accession** | **Probe sequence** | **HUGO gene** |
| --- | --- | --- | --- |
| PA0044 | NP_248734.1 | GCATATTCAATCATCTCAGCAGAACCCGTCTTTCG | exoT |
| PA0044 | NP_248734.1 | GCGACTTTGACAGCGGAGAGGCTGGCGAAGGATCA | exoT |
| PA0044 | NP_248734.1 | ACCTGAATCGGTCCCTGCGTCAGGGACGGGAGCTG | exoT |
| PA0085 | NP_248775.1 | AGGGTGAGTCCAAGGACAAGACTCACGCCGAGGAA | hcp1 |
| PA0085 | NP_248775.1 | TACCTGATCATCACCCTGAAGGAAGTCCTGGTGTC | hcp1 |
| PA0085 | NP_248775.1 | GGATGGCGCGAAGGACGGCGGTCCGGTCAAGTACG | hcp1 |
| PA0139 | NP_248829.1 | ACAACGGCAAGTTCATCGAGGTGACCGAGGAATCC | ahpC |
| PA0139 | NP_248829.1 | AGATCCACTCCAACGAGATCGCCCGTGACGTCGGC | ahpC |
| PA0139 | NP_248829.1 | ACGCCGCCAACAACTACGGCGAGTTCCAGAAAGCC | ahpC |
| PA0151 | NP_248841.1 | ACGTGGTGAGCAAGAAGCCACAACTGCAACAAGCC | PA0151 |
| PA0151 | NP_248841.1 | CTGCAACTGGCCGGGATGCAACATGACCTGCAATT | PA0151 |
| PA0151 | NP_248841.1 | CACCTACTACCCGTCCAGTGCCAATCGACTGTACG | PA0151 |
| PA0165 | NP_248855.1 | CACCGTGATGCTAGCCGGGGGGCTACTGGCCGCAG | PA0165 |
| PA0165 | NP_248855.1 | CGAAGACCATGACATGACGCCGACCCACGAGACCG | PA0165 |
| PA0165 | NP_248855.1 | GATCACTCCGCGCCTGAGCTTCGGCAAGCTCACCG | PA0165 |
| PA0192 | NP_248882.1 | ATCTCGGCGCGCTGAACGACACCTACTCGAAGGCC | PA0192 |
| PA0192 | NP_248882.1 | TCTCAGCCAGTACAACTTCGCCCCTTCGGCGTTGC | PA0192 |
| PA0192 | NP_248882.1 | TTGCGCGATCCGCAGCGCAGGAAGGTCAACATCAA | PA0192 |
| PA0423 | NP_249114.1 | CGGCTCCGCACTGTTCACCGCCGGCCAGGCAATGG | pasP |
| PA0423 | NP_249114.1 | TCGACGAGAAGAACCCGTCGGCCGACAAGGTCAAG | pasP |
| PA0423 | NP_249114.1 | TCGAATCCACCGAAGTGAAGGCCAACGGCGACAGC | pasP |
| PA0434 | NP_249125.1 | CGTTCCGGCCTTGCTCTGGGCCGATGAGGAACCCA | PA0434 |
| PA0434 | NP_249125.1 | GACGTGCGCAACCATCGGGTACCCGACAAGGACAA | PA0434 |
| PA0434 | NP_249125.1 | CCTACACGGTCTACGGGACTTTCCTCAGCTACCGG | PA0434 |
| PA0439 | NP_249130.1 | TGTCGTCGCGTTATTCCGCGCACTTCGGACCGAAC | PA0439 |
| PA0439 | NP_249130.1 | CGGCTGTCCGATTTCAGCGGCGGCGCGGTGGGCAA | PA0439 |
| PA0439 | NP_249130.1 | CTGCCCCGGGCCGACGGTACGCACCGCTACGAAGT | PA0439 |
| PA0470 | NP_249161.1 | GATATGAACAGCACCAGTTCGCTGAAGCCGACCGA | fiuA |
| PA0470 | NP_249161.1 | GCAGAACATAGGCGGCACGCCGGTCACTTCGCAGA | fiuA |
| PA0470 | NP_249161.1 | CCTATAGCTACGCCGACACCGAAGTGAAAAAAGGC | fiuA |
| PA0641 | NP_249332.1 | CTGCTGTTCGTCGAGTTCGATGCCAGCCAGTTCCA | PA0641 |
| PA0641 | NP_249332.1 | CAACGGTTACCAGAGCCAACCGGTAGCGGTGAACA | PA0641 |
| PA0641 | NP_249332.1 | GCCAGCAGTTCTTCTCCGATATCGAACGGATGCAG | PA0641 |
| PA0688 | NP_249379.1 | ACTTCCACCGTGGTGTCGACCGTCAAGGCAACCAA | PA0688 |
| PA0688 | NP_249379.1 | GCCTTCATCAACAAGCACTACGGTGGCACTACCAC | PA0688 |
| PA0688 | NP_249379.1 | CAGCGAAAACCTGGCCATCGGCAACACCAACGTCT | PA0688 |
| PA0690 | NP_249381.1 | CGGACAGATCACCTTGGATGGCATTCTGAACGCTT | PA0690 |
| PA0690 | NP_249381.1 | CAGAGCGATGTTGCCAGTGGATGCCAGATCCCTCT | PA0690 |
| PA0690 | NP_249381.1 | GGAAAGGCAAGACTGCATACGATTCGTCCGATGGT | PA0690 |
| PA0692 | NP_249383.1 | GCGTTCGCTGTTCAATTGTGCGCACCCGTTTCGCC | PA0692 |
| PA0692 | NP_249383.1 | CAGCATGAATGTGGTCGGCCAGGGCCACTCCTATG | PA0692 |
| PA0692 | NP_249383.1 | CTCCCAGGAATGGCGTACGCCGTCCCTCGGACGCT | PA0692 |
| PA0696 | NP_249387.1 | TGAACGCGAAAGTGAACCGACTGGCGCTGGGGATG | PA0696 |
| PA0696 | NP_249387.1 | GGGGAGGTCCGCGTACCCTATATCCCGCAGGTGGT | PA0696 |
| PA0696 | NP_249387.1 | GCCCTGGGCCGGAGATCCGGAGTGTGACACCGACG | PA0696 |
| PA0755 | NP_249446.1 | GCCTGGCAGCGAATGAACGGCGACGACGCCTTTCC | opdH |
| PA0755 | NP_249446.1 | ATGTCGACTTCCCTGCCGCAGGCCGCAAGGTCTGT | opdH |
| PA0755 | NP_249446.1 | CTGATTCCGAAGCTGCCATCGGTACAACCGAACAA | opdH |
| PA0781 | NP_249472.1 | GCCTTTTCCCTTCCCGTCCGCTCTGGCCTCTCACG | PA0781 |
| PA0781 | NP_249472.1 | GTGCCGGAAGACCCCAACAGCGTCGAGCGCCTGGA | PA0781 |
| PA0781 | NP_249472.1 | GCGCTGGCCTGCCTGATCGTTTCGGGGGAAACGCT | PA0781 |
| PA0844 | NP_249535.1 | AAAACTGGAAATTCCGCCGTCGAACCTTTCTCAAG | plcH |
| PA0844 | NP_249535.1 | TCTGGTACCAGAACTACAAGTACGAGTTCTCCCCC | plcH |
| PA0844 | NP_249535.1 | CAATCCGAACCGCCTCTACCACATGAGCGGACGCG | plcH |
| PA0852 | NP_249543.1 | ATGAAACACTACTCAGCCACCCTGGCACTCCTGCC | cbpD |
| PA0852 | NP_249543.1 | GCCAGCCGGTGCTACCGTCACCCTGCGTCTGTTCG | cbpD |
| PA0852 | NP_249543.1 | AAGGCCTGCAGCAATACGACGCCGGGACCGTAGTG | cbpD |
| PA0994 | NP_249685.1 | TAGCAACCGTACCTATGTGGAACGTGACATTCGTG | cupC3 |
| PA0994 | NP_249685.1 | CATGTCCGACGAGAGCTACCTCTATGGCTCAAGTT | cupC3 |
| PA0994 | NP_249685.1 | ACTCTGGAGGAAAGATTCCCTTCGGTGCACAGGCA | cupC3 |
| PA1092 | NP_249783.1 | TCGTGCCCGTACCGTGTTCACCGCTGATGTCAGCG | fliC |
| PA1092 | NP_249783.1 | ACCGGTGGTTCGCTGAACTTCGACGTAACCGTTGG | fliC |
| PA1092 | NP_249783.1 | ACTGCCAGCATCAACGACAAGGGTGTACTGACCAT | fliC |
| PA1248 | NP_249939.1 | AGTCCAAGTCCGGTTCGGAGAACACCTACAACCAG | aprF |
| PA1248 | NP_249939.1 | ACTACCTCACGGCCTGGGCACGCCTGCGTTTCTAC | aprF |
| PA1248 | NP_249939.1 | TGCACACCCTGTCGAAGACCGATACGGAGGAAAAC | aprF |
| PA1249 | NP_249940.1 | ACCTGACCTTCGGCAACTTCAGCAGTAGTGTCGGC | aprA |
| PA1249 | NP_249940.1 | CCAGCAATTCTCTTGCATTGAAAGGTCGTAGCGAT | aprA |
| PA1249 | NP_249940.1 | GCAGATCCTCCGCGAACAGGCGTCTTGGCAGAAAG | aprA |
| PA1302 | NP_249993.1 | GGTTTCACCTACCTGCGCTCGAAAAGCTCGCGCTG | PA1302 |
| PA1302 | NP_249993.1 | AAGCGGATGCGTACCGAGTACCAGGACAAGGTCCT | PA1302 |
| PA1302 | NP_249993.1 | GTCGATCGACAACGGCAGTATCGGTGGTAGCTGCG | PA1302 |
| PA1322 | NP_250013.1 | CCTCAAGCTCGAACACGATTTCAGCGACGACTTCC | PA1322 |
| PA1322 | NP_250013.1 | ATCCGCTATTCCCACCTGCATCGGGACACGGTGAT | PA1322 |
| PA1322 | NP_250013.1 | CCGCACCACCTACTCCTACAACCTGGGCAAGTTCT | PA1322 |
| PA1365 | NP_250056.1 | CGCAACAATCCTCGGTACAGATCCTTTTCGCCGGC | PA1365 |
| PA1365 | NP_250056.1 | CCGAAGAGGCATTGCGCCAGTTGCTCAGGGATAGC | PA1365 |
| PA1365 | NP_250056.1 | CGTGCCCGCTACGCAAGCCCGCAGCGAACCGCTGG | PA1365 |
| PA1383 | NP_250074.1 | CAAGAGTGTGGAAAATGGTGCCAACCTCGACAGCA | PA1383 |
| PA1383 | NP_250074.1 | CCAGCGTTACCTACACCATCGATCCAGTGAAGGCT | PA1383 |
| PA1383 | NP_250074.1 | CGTCCTGCATCCGTTGAACCTGAACAATCAGGATC | PA1383 |
| PA1493 | NP_250184.1 | CGGAGAAAGGCGAGAACATCACCATCCAGATGTCC | cysP |
| PA1493 | NP_250184.1 | CTCGCCGACAACGGCGGCCTGGTGCCGAAGGACTG | cysP |
| PA1493 | NP_250184.1 | TGAAGAACGGCGGTGACGAGAACAAGGCCAAGGAA | cysP |
| PA1494 | NP_250185.1 | GCCTGCAATTGCCCAAGGGCCAGCATGAGATCGCC | PA1494 |
| PA1494 | NP_250185.1 | CTACCCGCTGCTCCCCGGCACCCTGAATACCTTTC | PA1494 |
| PA1494 | NP_250185.1 | TCGGCAACCGCACGCGGGTCAACTACGAATTCCGC | PA1494 |
| PA1703 | NP_250394.1 | ACCCGGGTGTCTTCCGACGAATCGGCGGACCTCGG | pcrD |
| PA1703 | NP_250394.1 | CGTTGTACCTGGATTTCGGCGTGCCGTTCCCGGGT | pcrD |
| PA1703 | NP_250394.1 | CGATGGCGAATACCTGGTGCAGCTACAGGAGGTCC | pcrD |
| PA1708 | NP_250399.1 | GGTCGCGAGGGATGTCGAGAGTCTGCGGGTGGAGC | popB |
| PA1708 | NP_250399.1 | TCTGGCAGCCAAAATCTTTGGTTGGATCAGTGCAA | popB |
| PA1708 | NP_250399.1 | GGGCGGTGTCATGGGAGTGGTCTCCCAGTCGGTAC | popB |
| PA1709 | NP_250400.1 | CGACACGCAATATTCCCTGGCGGCTACCCAGGCCG | popD |
| PA1709 | NP_250400.1 | GCAAGGCCATCAGTCAAGAGAAGACCCTGCAGAAG | popD |
| PA1709 | NP_250400.1 | ATGAAGGACGTCCTGCAGCTCATCCAGCAGTACAC | popD |
| PA1764 | NP_250455.1 | GTGGGTTTCTCCCACCAGGCAGTAGCGTTGAACGA | PA1764 |
| PA1764 | NP_250455.1 | TCGCCAAGTACCTCGACTCGAAGGACGCCACCAGC | PA1764 |
| PA1764 | NP_250455.1 | CTATCCGGATGCCTTCTCCACCGGCGTGAAGCTGC | PA1764 |
| PA1804 | NP_250495.1 | TTGCCGGTCGCGCACTGGACGCAGTGATCGAGTCC | hupB |
| PA1804 | NP_250495.1 | GACTCCGTCGTGCTGGTTGGTTTCGGCACCTTCGC | hupB |
| PA1804 | NP_250495.1 | CGCAAACTGGCAAGCCGATCAAGATCGCCGCTGCC | hupB |
| PA1910 | NP_250600.1 | ACCATCAGAGCCGCGAGTTTCCCATGCTCTCCCTG | femA |
| PA1910 | NP_250600.1 | TCGAGATCAGGAAACCGAACGCCTATACCGACGCC | femA |
| PA1910 | NP_250600.1 | GTCGATCAGCTCGCCGCGCACCCTTTCCCTTTCCC | femA |
| PA1974 | NP_250664.1 | AGGTCGACGAGCTGAGCCTCGGTGGTCGCGCTTCC | PA1974 |
| PA1974 | NP_250664.1 | CCGCGGCGCATTGCTGCAGGGCTCGTATTCATGGG | PA1974 |
| PA1974 | NP_250664.1 | AACACCGCCGTTCGCCGAATCCTGCCGCTGACCCT | PA1974 |
| PA2019 | NP_250709.1 | CCAAAGGGCTTGGTGGAAGACGTGGAGGTCCGCGC | PA2019 |
| PA2019 | NP_250709.1 | CCTGCGCCCTTGAAGGCGGCCCTGGACATCAGCCG | PA2019 |
| PA2019 | NP_250709.1 | GCCCTATTCCTGCTGGGCTGCGAAGAAGCAGCGGA | PA2019 |
| PA2057 | NP_250747.1 | AACGTCTTCGAGGCCTTGCAGACGCAGACCCAGAA | PA2057 |
| PA2057 | NP_250747.1 | GTGTTGTCCCACGACTACAAGCAGTCCGACGACTA | PA2057 |
| PA2057 | NP_250747.1 | ACCTGGGACTACGCACAGTTCACCTCGACGCTCTT | PA2057 |
| PA2070 | NP_250760.1 | GGCTTTCCGGCAACCAGGGCGTATACAACTCCAGT | PA2070 |
| PA2070 | NP_250760.1 | CCTCGTACAACCAGAACAAGGTGGTCGACCATATC | PA2070 |
| PA2070 | NP_250760.1 | CGCAAGGAAGCGTTCCACCAGGACATCCAGGACTT | PA2070 |
| PA2089 | NP_250779.1 | ATCACCAGCGACGAGGACGACTATTCGCTGACCCT | PA2089 |
| PA2089 | NP_250779.1 | GAGATCGGCATGGACCATTCGATCAACCTCGCCCT | PA2089 |
| PA2089 | NP_250779.1 | CGCCGAAGACCTGGAGCCGGAAAAATCGATCAGCT | PA2089 |
| PA2128 | NP_250818.1 | TTCGAACCCATGCGCAGTGGTATTGGCCTTTGCGG | cupA1 |
| PA2128 | NP_250818.1 | GGCGGCAAACACTATCACATTCAGCGGCGAAGTGA | cupA1 |
| PA2128 | NP_250818.1 | TTCACCCTGCAACTCACCGATTGCGTCGCGCCGAC | cupA1 |
| PA2204 | NP_250894.1 | CGGAAGTCCTCAACAAGTGGCGGGTCGGCGTGGAC | PA2204 |
| PA2204 | NP_250894.1 | CCCGACCGCGACAAGTACGAAGTGCCGCCCTTCAC | PA2204 |
| PA2204 | NP_250894.1 | CTGATCGGCGTCGGTATTCCCAAGGGCGAGAAGGC | PA2204 |
| PA2289 | NP_250979.1 | AGCGCAGTGTCGTGCAGTATCAGGCGATACCCAAG | PA2289 |
| PA2289 | NP_250979.1 | GCCAGAGAAACGGCGCGGTGATCGACTTCGACCGC | PA2289 |
| PA2289 | NP_250979.1 | GGGCTGTCCTGGTACGCGGGAGCCGGGGTCGAATA | PA2289 |
| PA2291 | NP_250981.1 | AAGTACAACTTCGCGCCGGACTGGTACGTGCAGGT | PA2291 |
| PA2291 | NP_250981.1 | GCGCTGGTGGCGGGGATCAAGATCCAGACGGTGTT | PA2291 |
| PA2291 | NP_250981.1 | CACGCCGCCGAGGCGTTCTCGCCGAACTCGAAATG | PA2291 |
| PA2296 | NP_250986.1 | GCCCTGACTTCCCTGGGTGCCCAGGCGGAAACCAT | PA2296 |
| PA2296 | NP_250986.1 | AGGAGTTCGAACGGGTAGCGCAGATCTGGGTGAAG | PA2296 |
| PA2296 | NP_250986.1 | AGGGCAAGGCGATTCGTGCGATCTACGCCCAGGAT | PA2296 |
| PA2367 | NP_251057.1 | ATCAAAGGCGACAGCCTGCTCGTCGGCTACGAAAA | PA2367 |
| PA2367 | NP_251057.1 | CCCACGTCGGCGAATTCACCCTCACCAAGTTCATC | PA2367 |
| PA2367 | NP_251057.1 | GCCGGCAAGCCGATCGCCGAGGCGACCATCACCAT | PA2367 |
| PA2420 | NP_251110.1 | GCTCTGCCTTGCCCTTGCCATAGGTCCGGGCTCGA | PA2420 |
| PA2420 | NP_251110.1 | GGCGTCAGTCGCATGGTGCCGCAGAGCTACCGTGG | PA2420 |
| PA2420 | NP_251110.1 | CCCTGAAGAACTACCATTTCCGCGCGCTCGAACTG | PA2420 |
| PA2505 | NP_251195.1 | ACCAGCGGCAGCGACGACTTCTACAGCTTCTACAC | opdT |
| PA2505 | NP_251195.1 | GGAGTGGGTGCAGGGCTTCATGGCCAACTTTTCCT | opdT |
| PA2505 | NP_251195.1 | ATGACGACCAGGGGCGGCCGCAGGACGACTATTCG | opdT |
| PA2590 | NP_251280.1 | GATGTACGTCTTTCCTCTCGTTCGTTGACCCCCGT | PA2590 |
| PA2590 | NP_251280.1 | TCGCCACGAAACGGCCAATCTCAATATCAGCGGGG | PA2590 |
| PA2590 | NP_251280.1 | CACCGGCAATCGTTGGCGCTCAACTCGGGTTTCGG | PA2590 |
| PA2662 | NP_251352.1 | GCATATTTCTGGATCGCCGTGGCGCCGCTTGGCAT | PA2662 |
| PA2662 | NP_251352.1 | TGCCCCTGGCAGGCGGACTGTGGGCTTTGGCGTTC | PA2662 |
| PA2662 | NP_251352.1 | CTGGCATATTTCTGGATCGCCGTGGCGCCGCTTGG | PA2662 |
| PA2688 | NP_251378.1 | GCCGGCGCGGAAACCCACGGTAATCTCAGCGTCTA | pfeA |
| PA2688 | NP_251378.1 | AAGGTATCTTCGACCCCAACAACGCCGGCTTCTAC | pfeA |
| PA2688 | NP_251378.1 | CTCGATCCCCGGTCTCGCCGGAAAGAACCGCAGCA | pfeA |
| PA2862 | NP_251552.1 | TCGGTCTCGCCTCTCTCGCTGCCAGCCCTCTGATC | lipA |
| PA2862 | NP_251552.1 | CACCGGTACGCAGAATTCACTGGGCTCGCTGGAGT | lipA |
| PA2862 | NP_251552.1 | TTCCTCGGCGCCTCGTCGCTGACCTTCAAGAACGG | lipA |
| PA2911 | NP_251601.1 | ACTCGACCACATCGACCTCGCCACTCCGGTCAGTG | PA2911 |
| PA2911 | NP_251601.1 | CCGCTTCAGCCGTTTGCGCACCGAGCCTTTCGCCC | PA2911 |
| PA2911 | NP_251601.1 | CCTGGCTGACAACAACAACGGCCAGCAATGGATCG | PA2911 |
| PA2939 | NP_251629.1 | CCGCATCCCTGGCGGCACCTTCGGAAGCGCAACAG | PA2939 |
| PA2939 | NP_251629.1 | CGCCTACTACCCGAAAGGCCCGGGTAGCCTGAGCG | PA2939 |
| PA2939 | NP_251629.1 | GCCAGAAAGCACAATCGCGGTCGCTGCAGATGCAG | PA2939 |
| PA3105 | NP_251795.1 | CCGGAGGGACGACCCAGCATCCTGCCGACCAACGC | xcpQ |
| PA3105 | NP_251795.1 | GAGCAACAAGGCACCGGAATCCATCCCCGACGGCG | xcpQ |
| PA3105 | NP_251795.1 | GCGCAACAATACCGACCTGATCACCAGCAAGCGCT | xcpQ |
| PA3162 | NP_251852.1 | AAGAACTGCGTAAGCAGGAAGTAGAAAGCGCTGGT | rpsA |
| PA3162 | NP_251852.1 | GTCATCAGCCTGGGCGACGACATCGAAGGTATCCT | rpsA |
| PA3162 | NP_251852.1 | GCGTCCCTGCACGAGAAAGGCAGCATCGTCCGCGG | rpsA |
| PA3268 | NP_251958.1 | CAGCGTGGTGCGTCGTCGGGAAATGCTCGAGAGCG | PA3268 |
| PA3268 | NP_251958.1 | CGCAAGGACTTCTCGCTGAAGTACACCCGCCAGGT | PA3268 |
| PA3268 | NP_251958.1 | CGACAGCCAGGGCAACTTCACCCACAACTACATCA | PA3268 |
| PA3280 | NP_251970.1 | CAACCGCGCGACCTTCGGCGGTGTCTCCAACTCGC | oprO |
| PA3280 | NP_251970.1 | CAGTACAAGCTGGAAGGTGCCAAGTTCGACTCCGT | oprO |
| PA3280 | NP_251970.1 | GGGCGCCCAGGTCAACTCGACCCTCGCCGACATGG | oprO |
| PA3422 | NP_252112.1 | AAAAAGGCGCGGAATCTTCCCTCTGCACGCACTCG | PA3422 |
| PA3422 | NP_252112.1 | GCTCAACGTGGCCAACCCGCAGGAGTGCAACAGCC | PA3422 |
| PA3422 | NP_252112.1 | GTGGGATCCTCGAATATCCCAACGTGTTCGCCGGG | PA3422 |
| PA3526 | NP_252216.1 | GCGCCTTCTTCTATTGCCTTTCCTGCTCAGCAGCC | PA3526 |
| PA3526 | NP_252216.1 | AAGAGCAACGGGGTGCCGGAAAGCCAGATCAACGT | PA3526 |
| PA3526 | NP_252216.1 | CCTCCGGCCGCCTCGCTTCAGGGAAAACCCACGGT | PA3526 |
| PA3544 | NP_252234.1 | ACCCTGCAATCGGACACCGACGACGGCAACAACAG | algE |
| PA3544 | NP_252234.1 | CCACCCAGTGGGCGCCGCACCACCGCATAGGCGTG | algE |
| PA3544 | NP_252234.1 | TACCGGCGATGCCTACAACTATCGTTCGAGCATGC | algE |
| PA3724 | NP_252413.1 | GCGCATTTCTTCGGCGGCGTGGTGTTCAAACTGTA | lasB |
| PA3724 | NP_252413.1 | AACTCCGGGCTGATCTACCGCGGGCAATCAGGCGG | lasB |
| PA3724 | NP_252413.1 | TGATCGACGTGTCCAAACTCCCCAGCAAGGCTGCC | lasB |
| PA3790 | NP_252479.1 | GCGAGCTGGGCGCTTCCTACCAGCTCACCGGCAAC | oprC |
| PA3790 | NP_252479.1 | CTTTCAGCTTCGACCCTGCCCGTCTCGCACAACGC | oprC |
| PA3790 | NP_252479.1 | TCCAAGTACGACATGATGACCGACTACTACACCGA | oprC |
| PA3841 | NP_252530.1 | ATCGAACTACAAGAATGAAAAAGAGATTCTCTATA | exoS |
| PA3841 | NP_252530.1 | CGATCATGGACTGGCTGGGCAAGCTGTTGGGCTCC | exoS |
| PA3841 | NP_252530.1 | TACCTGGGCAGACAGCCTGGTGGCATCCACAGTGA | exoS |
| PA3842 | NP_252531.1 | CGCGCCGCCATCCACCAGCTTTTTCTCGCTCTCGA | PA3842 |
| PA3842 | NP_252531.1 | TGGCAGAACATCCGACCGATCACCTGTTGATGTTC | PA3842 |
| PA3842 | NP_252531.1 | CTACGGCCAACGAGCAGAACCTGTTCAGCCAGGAC | PA3842 |
| PA3866 | NP_252555.1 | GAAATGATAATGGTTACGAGCCTGGGTCGATGGTT | PA3866 |
| PA3866 | NP_252555.1 | GTGGTAAATCGCAGATTGCCGCGATGTTGACCAAC | PA3866 |
| PA3866 | NP_252555.1 | CAAATTGCAGATCGGAACGATTCCGCATGGGCGGT | PA3866 |
| PA3901 | NP_252590.1 | GATCGATTCCGAACAGAAGAACCTGCTGAAGAACA | fecA |
| PA3901 | NP_252590.1 | AACTCAGCCCCAGCTCCAGCCAGGACGGCCTGAAG | fecA |
| PA3901 | NP_252590.1 | AGCGACGAACACACGTTTTCGGCGATGACCCAGTA | fecA |
| PA3923 | NP_252612.1 | ACCCGTAATCCGTTCTCGCGTCGGGCCCGGTTGCC | PA3923 |
| PA3923 | NP_252612.1 | TTACCCCGTCGCCCTGGGACGAGGGCATCCTGGTC | PA3923 |
| PA3923 | NP_252612.1 | ACCATCTCCGGCCTGGCCAGCTGCCAGGCGCTGAA | PA3923 |
| PA4082 | NP_252771.1 | AGTTCAACGGCAACTCCTCCAGATACCGTTTCACG | cupB5 |
| PA4082 | NP_252771.1 | GTATCCGATAACACCTACGGCTCCGGACCGTCTTC | cupB5 |
| PA4082 | NP_252771.1 | CGTATCGTTTCGTCCATGTCGGAAGGCACCGTCAG | cupB5 |
| PA4103 | NP_252792.1 | CCCAACCTCTCAATCCGCCTGGCACGGCTGGCGAC | PA4103 |
| PA4103 | NP_252792.1 | GCAGTACACGCCTTGCTGATCCTGGCCTACTGGAA | PA4103 |
| PA4103 | NP_252792.1 | CCCCGAGACATTCTGGAGCGGCCGCTCGGTCGCGG | PA4103 |
| PA4107 | NP_252796.1 | GCGGAGAAGGCAAGTGTGGTAGCGGCGGCTCCGCG | PA4107 |
| PA4107 | NP_252796.1 | TCGATAGCGACCATGACGGCTTCATTTCCGAAGCC | PA4107 |
| PA4107 | NP_252796.1 | CGTACTGGTGGGGGGAATGCTGCTCGGCGGTTCGG | PA4107 |
| PA4156 | NP_252845.1 | AACGCCTCGGTCGCCGAGGTGATCAATGGCAGTCC | PA4156 |
| PA4156 | NP_252845.1 | TCGGCCTTCCTGCCGAAAGTCTCCCTGGCCTTCGC | PA4156 |
| PA4156 | NP_252845.1 | GCCAACACCAAGGCCTACTCGATCAAGGCCTACAC | PA4156 |
| PA4168 | NP_252857.1 | ACGTCGTTTCCAGTGCGTTATTGCTCGGTCTCTCC | fpvB |
| PA4168 | NP_252857.1 | CAGGTTCCGGACTGAGCGGGGAGGTTCGGGGCGGC | fpvB |
| PA4168 | NP_252857.1 | CAGAAAGGCAACAGCTTCCAGGACCACGTCAGCAG | fpvB |
| PA4175 | NP_252864.1 | ATAAGAGAACGTACCTGAATGCATGCCTGGTTCTG | piv |
| PA4175 | NP_252864.1 | GCGCCCACCTCACAGCGCAATGACTACTTCTCCGA | piv |
| PA4175 | NP_252864.1 | GGTCAACCGTCCCTACTGGAGCCCGGTCATCGAGG | piv |
| PA4195 | NP_252884.1 | GCAAGGCGTTGCCCAACGCCCAGGTGGTCACCTTC | PA4195 |
| PA4195 | NP_252884.1 | GCAATCCGCAACTCAAGCAACTGGTCGACTCGACC | PA4195 |
| PA4195 | NP_252884.1 | AGCGCCACTTCAAGATCGATTCCGACCGGATCCCC | PA4195 |
| PA4213 | NP_250593.1 | TATACGCCCACGTCGGGGTGCTGATCTCCACGGTC | phzD2 |
| PA4213 | NP_250593.1 | AGGACTTCTGGGGGCCGGGCATGCGCGCCAGTCCG | phzD2 |
| PA4213 | NP_250593.1 | CGAAATCACCGCCTATCCCTTGCCGACCGCCCAGC | phzD2 |
| PA4221 | NP_252911.1 | AGGCACTGGCTACCTGACGCCCTACTGGAGTCTCT | fptA |
| PA4221 | NP_252911.1 | CAAGGACAGCCAGAACGACTCGGGGACGCGCTACT | fptA |
| PA4221 | NP_252911.1 | AAAACGGAGACGAAGGTGATCAAGGGACGCCAGGG | fptA |
| PA4271 | NP_252961.1 | GGCCGTCGTAGACGGCGCTCCGGGCGTGGTGAAGG | rplL |
| PA4271 | NP_252961.1 | CATCGTCCTGGCCGAAGCTGGCGACAAGAAAGTGA | rplL |
| PA4271 | NP_252961.1 | TTCGGCGTTACCGCTGCTGCCGCTACCGTTGCCGC | rplL |
| PA4273 | NP_252963.1 | CGAGAAAGTAGTTGCCGGCAAGCAGTACAGCTTCG | rplA |
| PA4273 | NP_252963.1 | TCATCCACAGCTCCGTCGGTAAGGTCGATTTCGAG | rplA |
| PA4273 | NP_252963.1 | GCCTGAAGCCGTCCTCGTCGAAAGGCGTGTACGTC | rplA |
| PA4277 | NP_252967.1 | AGCGCCGTGCAGAAGCTGGTAGAGACCCTGGACTC | tufB |
| PA4277 | NP_252967.1 | GTCCAGGAAGAAGTGGAAATCGTCGGCATCAAGGC | tufB |
| PA4277 | NP_252967.1 | CTGCACTGACCAAGGTCTGCTCCGACACCTGGGGT | tufB |
| PA4370 | NP_253060.1 | TCAAGAATTACGCGGACCTCGCCGAAGCCACCTTC | icmP |
| PA4370 | NP_253060.1 | CAAGGAAGCCTGGTTCGCCGCTCGTACCCCTTACT | icmP |
| PA4370 | NP_253060.1 | GCAACCGTCCCGCCACCGACTATGCCCAGGGCAAG | icmP |
| PA4385 | NP_253075.1 | CGTGGCTATCGTCGCCCAGCTGAAAGAGCTGGCCA | groEL |
| PA4385 | NP_253075.1 | CACCACCATCATCGATGGCGCCGGTGTGCAGGCTG | groEL |
| PA4385 | NP_253075.1 | AGCCCTGGTTCGTGCCCTGCAAGCCATCGAAGGCC | groEL |
| PA4514 | NP_253204.1 | CTGTCCTATTACCATCTCAGCACCGATGATATGCC | PA4514 |
| PA4514 | NP_253204.1 | TGAAGCTAAGCGAACAATGGGAGCTGAATCTCGGT | PA4514 |
| PA4514 | NP_253204.1 | GTGGACCACCTACGATCTGTTGCAGAACTTCACTA | PA4514 |
| PA4525 | NP_253215.1 | TCGCGTGGAATTGCTGGTAGCAAAATTAAAATTGG | pilA |
| PA4525 | NP_253215.1 | GTAGCAATCGAAGATAGTGGTGCGGGTGATATTAC | pilA |
| PA4525 | NP_253215.1 | TTGTAAATCTACCCAGGATCCGATGTTCACTCCGA | pilA |
| PA4554 | NP_253244.1 | AGCATTTCCGGCTATGGAAACTATACGTTCTTCGC | pilY1 |
| PA4554 | NP_253244.1 | TTCTTCAATTGGCTGGAAAAACTTTCGGTCAATGG | pilY1 |
| PA4554 | NP_253244.1 | CCGGATATCGACGACAATATCAAACCGTACATTCC | pilY1 |
| PA4589 | NP_253279.1 | CCGCAACAAGCTGAAGGGGCACTACAACCTCTACG | PA4589 |
| PA4589 | NP_253279.1 | TGGATGGCAGGGCAACCAGCAAGGGACAGGTGTTC | PA4589 |
| PA4589 | NP_253279.1 | AGGCGGCTCGCTGTTTCCCAACGATCCGAGCGCCG | PA4589 |
| PA4624 | NP_253314.1 | AACGTCCGCTGGATGCCGATACCCTGGAGCGCGGC | PA4624 |
| PA4624 | NP_253314.1 | CCTTGCAACAGCAGCGATACGAAGGCTTCGTCAGC | PA4624 |
| PA4624 | NP_253314.1 | GCAACATCGGGCGTCTGCAAGGGCTCGGCGGACCC | PA4624 |
| PA4709 | NP_253397.1 | AAGTGACGGTCTCTGCCAATGGGCAGATGGGCCTG | PA4709 |
| PA4709 | NP_253397.1 | TGGATAATGGCGACCTGGCGAAACTCTTCGAAGCG | PA4709 |
| PA4709 | NP_253397.1 | TTCGCCATTGCCGAGGAGACAGCGCGGGGTACCCA | PA4709 |
| PA4710 | NP_253398.1 | GACTTCATCGACGAAGACGCCCTGAATACCGATAG | phuR |
| PA4710 | NP_253398.1 | CCGTACCGAGCAGGCAGTGGATTCGGTGCCAAGCA | phuR |
| PA4710 | NP_253398.1 | CTGCAGGCGGGCTACCACATCGAGCCTAACCCCAA | phuR |
| PA4837 | NP_253524.1 | ACAACAACGAAACCCTCCAGGCCACCCTGCGCCAT | PA4837 |
| PA4837 | NP_253524.1 | CCTGAACAATGGCCGCTTCCGCGAAACCAGCTCGC | PA4837 |
| PA4837 | NP_253524.1 | ACGCCTCGAAATCCTTCAAGCCCAATGGCGGTACC | PA4837 |
| PA4881 | NP_253568.1 | TGAAAGCTTCCCTGATCCTCGGCCTCGCCCTCGCC | PA4881 |
| PA4881 | NP_253568.1 | ACCGGCTGCACAAGGATATGTCCAGCGTCGAACTG | PA4881 |
| PA4881 | NP_253568.1 | GCCTGCATCGCAGCATGAGCCGTTTCGACCTGCGC | PA4881 |
| PA4897 | NP_253584.1 | GCCAACCCTGCGAGCGAAGAAGTCATGTCCCTACA | PA4897 |
| PA4897 | NP_253584.1 | TCGCGAGGAAAGCAGCGTAGCCAAGGTCTACAACG | PA4897 |
| PA4897 | NP_253584.1 | TGGTCACCGGCTACAACTACAGCAAGAAGGGCAGC | PA4897 |
| PA4898 | NP_253585.1 | AATCCGTCGTAAAACCGCAGTCCCGCGCAATGCAG | opdK |
| PA4898 | NP_253585.1 | CCAGCAATTCCGAGCTTCTTCCGTTGCACGATGAC | opdK |
| PA4898 | NP_253585.1 | TCGACAGCCATACCGTCTACGGCCTGTTTTCCGCC | opdK |
| PA4922 | NP_253609.1 | TGCTACGTAAACTCGCTGCGGTATCCCTGCTGTCC | azu |
| PA4922 | NP_253609.1 | CCTGCTCAGTGCGCCACTGCTGGCTGCCGAGTGCT | azu |
| PA4922 | NP_253609.1 | GCAGTTCACCGTCAACCTGTCCCACCCCGGCAACC | azu |
| PA5089 | NP_253776.1 | GGCTGACCCTCGACTGGTTCGCCAACAAGGCCTTC | PA5089 |
| PA5089 | NP_253776.1 | AGCAGGTCAGGGAAAACCTACATCCCGATCCCTAT | PA5089 |
| PA5089 | NP_253776.1 | GAGGAGTTGTGGGAGCTACATGCCCAGAAAGTGGC | PA5089 |
| PA5171 | NP_253858.1 | AAGTTCCACCCCGAGTTCGCCAACGCCGAGTTCGA | arcA |
| PA5171 | NP_253858.1 | GGCATGAACATCCGCCGCGAGGAGAAAACCTTCCT | arcA |
| PA5171 | NP_253858.1 | GACGACCTGCCCGCCAGCGAAGGCGCCAACATCCT | arcA |
| PA5340 | NP_254027.1 | TGACGAGCACTGGAAGGTTGCCTTCTATGGCGACG | PA5340 |
| PA5340 | NP_254027.1 | GCGGTTTCGAGCGGGCCTTGTATGGCAGCTATCTG | PA5340 |
| PA5340 | NP_254027.1 | CAACAAGGCAACCGCGGCCGCGAGTTGCTGTTCAG | PA5340 |
| PA5370 | NP_254057.1 | GAATGCTCCCGAGGCGCCGGCCCAGCGTCCATTGA | PA5370 |
| PA5370 | NP_254057.1 | TGGCGGATCGTTACGGCGCGGGGCTGACTTTCGTT | PA5370 |
| PA5370 | NP_254057.1 | GCTTACTACGTGGTCACTCTGTGCCTGGTCGGCTT | PA5370 |
| PA5464 | NP_254151.1 | CAGCAACTCTTGGACGAAGCCGCGGACAAGTTGCC | PA5464 |
| PA5464 | NP_254151.1 | GGCGCCGGAACACAGTGCCTGTGCGCAGAGCTTCG | PA5464 |
| PA5464 | NP_254151.1 | GAGTTGAAGCGCCTGTACCTCGGCCATCTGGACAA | PA5464 |
| PA5501 | NP_254188.1 | CGAAGGTGGAAGGCCTGCCGGTCGGAGCGATCCGC | znuB |
| PA5501 | NP_254188.1 | GCTGGCGGTGAGCGCCAGCGACCTGCTGTGGATAA | znuB |
| PA5501 | NP_254188.1 | ATGGCGGTCGGTGCCAGCCTGCTGGGACTGGTCGC | znuB |
